# Low-Frequency activity shapes fine-scale information routing in the early visual cortex

**DOI:** 10.64898/2026.05.25.727722

**Authors:** Danila Shelepenkov, Gregory Acacia, Mathilde Bonnefond

## Abstract

Visual processing requires flexible routing of task-relevant information across cortical hierarchies. One proposed mechanism is nested oscillatory activity, in which low-frequency rhythms dynamically modulate local excitability and inter-areal communication. However, the specific predictions of this framework have not been tested during active stimulus processing at fine spatial and temporal scales. Here, we reanalyzed local field potentials (LFPs) and multi-unit activity (MUA) recorded from V1 and V4 in macaque monkeys performing a figure–ground segregation task. We show that alpha-band activity in V1 carries information about the position of the figure and the orientation of the stimulus with fine spatial specificity within a transient post-stimulus window, a period that coincides with the emergence of figure-ground modulation and the dominant V4*→*V1 feedback. During this window, both local spiking activity and inter-areal coupling between V1 and V4 depended on alpha amplitude and on the instantaneous V1–V4 phase difference, indicating that low-frequency synchronization shapes effective communication between cortical populations. Together, these findings support the view that alpha-band activity reflect fine-scale information routing during visual processing through coordinated modulation of local excitability and hierarchical feedback interactions.

## 1 Introduction

The brain continuously processes an ongoing stream of sensory input, requiring efficient and flexible allocation of limited computational resources to the most relevant information. This requires dynamic mechanisms capable of selectively suppressing irrelevant signals while enhancing relevant ones. Oscillatory activity has been proposed as one of the leading candidates for implementing such flexible routing in neuronal populations (Buzsáki & Draguhn, 2004; Hoppensteadt & Izhikevich, 1998; Singer, 2018). Neuronal oscillations reflect coordinated fluctuations in population-level excitability, possibly allowing the temporal organization of neural communication across distributed areas of the brain. Empirical and computational work suggests that these dynamics improve computational efficiency and may implement routing functions by selectively gating inter-areal communication (Singer & Effenberger, 2025).

Several theoretical frameworks have been proposed to explain how oscillations may support information routing. The communication-through-coherence hypothesis proposes that inter-areal communication is facilitated by stable phase relationships between neuronal populations, particularly in the gamma band (> 30 *Hz*), allowing precise temporal alignment of spiking activity across regions (Fries, 2005). This idea was later extended to include a role for theta activity (4 to 7 *Hz*) in controlling gamma (Bastos et al., 2015; Fries, 2015) and further integrated into predictive routing accounts, in which oscillatory phase relationships determine which signals are routed forward or suppressed within cortical hierarchies (Gabhart et al., 2025). Instead of focusing on phase synchronization, the gating-by-inhibition hypothesis suggests that routing is achieved by functionally inhibiting irrelevant regions through an increase in alpha-band power (8–13 Hz) (Jensen & Mazaheri, 2010), reflecting inhibitory pulses that occur approximately every 100 ms (Klimesch et al., 2007). However, the precise role of alpha in actively suppressing irrelevant processing remains debated (Clayton et al., 2018; Jensen & Bonnefond, 2026).

Bonnefond et al. (2017) proposed a unifying framework in whic both mechanisms operate together: feedforward spikes and high-frequency activity that carry stimulus information are nested within low-frequency feedback cycles that rhythmically gate transmission. In task-relevant populations, selective routing is achieved by a decrease in low-frequency amplitude combined with inter-areal phase synchronization, while irrelevant populations are concurrently suppressed by an amplitude increase. This framework generates several specific, testable predictions. First, low-frequency oscillations must exhibit fine-scale spatial specificity, that is, receptive field or feature selectivity, consistent with their proposed role in routing task-relevant signals. Second, spikes and high-frequency activity should encode stimulus-specific information and be modulated by the amplitude and phase of local low-frequency oscillations. Third, inter-areal phase synchrony of low frequencies should determine the amount of information transmitted via spikes and high-frequency activity.

Although some evidence supports individual elements of this framework, including spatial specificity of alpha activity (Clausner et al., 2025; Harvey & Dumoulin, 2013; Yuasa et al., 2025) and modulation of gamma and MUA by alpha in V1 (Bollimunta et al., 2008, 2011; Bonnefond & Jensen, 2015; Dougherty et al., 2017; Spaak et al., 2012), the different predictions of the framework have not been tested together during stimulus processing.

To address this gap, we reanalyzed an existing dataset of local field potentials (LFP) and multi-unit activity (MUA) recorded from visual areas V1 and V4 in two macaque monkeys performing a figure–ground segregation task (Poort et al., 2012; van Kerkoerle et al., 2014). The task manipulated the position of the figure relative to the receptive field (RF) of the recorded channels, allowing us to investigate the fine-scale spatial modulation in V1 depending on the higher-order cortical processing in V4. Importantly, previous analyses of this dataset have already established several relevant findings: the amplitude of gamma oscillations and MUA in V1 and V4 increases when stimuli are presented in the figure condition compared to the background, whereas low-frequency power tends to be stronger in background conditions. In addition, V4 exhibits an alpha phase lead relative to V1, suggesting that activity in the alpha band reflects feedback processing. Finally, microstimulation in V1 elicited gamma oscillations in V4, while microstimulation in V4 elicited alpha oscillations in V1 (van Kerkoerle et al., 2014). These results provide a strong empirical foundation for testing the framework’s predictions in this dataset.

Moreover, figure–ground segregation is theoretically well characterized process in terms of its temporal dynamics, making it possible to link low frequency modulations to specific processing stages. Following the initial visual response, an early edge detection phase emerges, largely independent of attention and likely arising from a bipartite specialization of neurons in V1 and local horizontal connections (Ding et al., 2026; Westerberg & Roelfsema, 2025). However, the presence of border-ownership signals in V4 indicates that hierarchical processing may also contribute to boundary assignment (Jeurissen et al., 2024). Edge detection is followed by figure filling and background suppression steps, both of which depend on top-down attentional feedback (Kirchberger et al., 2021; Klink et al., 2017; Lamme et al., 1998; Markov et al., 2011; Poort et al., 2012, 2016; Self et al., 2019; Self et al., 2013; Supèr et al., 2001). These later processes have been proposed to be temporally dissociable and independently modulated (Poort et al., 2016), and possibly differentiated into subprocesses (Self et al., 2019) although this distinction remains debated (Jeurissen et al., 2024) and is not directly testable in our dataset.

Based on these findings and the framework outlined above, we derive the following predictions for the figure–ground segregation task.

We hypothesize that successful figure processing is associated with a spatially specific alpha amplitude over figure positions, driven by top-down interaction between V1 and V4. We predict that this reduction at figure positions, together with a corresponding increase at background positions, emerges after early edge detection, beginning around 90 ms after stimulus onset and extending to approximately 250–400 ms, consistent with behavioral response times in similar tasks (Poort et al., 2012). If border-ownership cells in V4 contribute to early modulation as proposed by Jeurissen et al. (2024), an earlier alpha modulation coinciding with the onset of MUA edge responses may also be observed at edge positions. These predictions imply that figure position can be decoded at fine spatial scale from alpha amplitude and that MUA and alpha amplitude are negatively correlated.

Second, we predict that V1 MUA will be stronger at a specific alpha phase during the time window associated with figure selection. We further predict that both alpha-amplitude modulation and the instantaneous alpha-band phase difference between V1 and V4 influence information transfer between the two cortical areas.

Third, beyond spatial selectivity, we predict that alpha activity will show feature-specific resolution, as recently demonstrated by Clausner et al. (2025). Specifically, figure features should be decodable from alpha amplitude within the same temporal window as position decoding, with decoding weights opposite in sign to those obtained from MUA, consistent with the negative relationship between MUA and alpha activity. Finally, we predict that alpha frequency will be higher in electrode groups exhibiting feature-selective responses, in line with the proposal that increased alpha frequency reflects a higher sampling rate for task-relevant features.

## 2 Results

To test our hypotheses we reanalyzed local field potentials (LFP) and multi-unit activity (MUA) recorded in visual areas V1 and V4 of two macaque monkeys performing a figure-detection task (Poort et al., 2012). The positions of the figure were systematically varied along the horizontal axis in fixed steps of 0.5°, ranging from 17 to 23 positions depending on the animal (see Figure 4). This design allowed us to test our prediction that figure position can be decoded in fine scale from alpha-band activity following the initial feedforward response.

### Decoding figure position

To avoid confounding orientation-dependent neural responses with figure position relative to the receptive field (RF), we generated stimulus combinations in which figure and background gratings shared identical orientations; edge locations still contained both orientations in both conditions. The position of the figure was quantified as the distance from the RF center of the recorded channels. The decoding was performed using ridge regression with two spatial profiles: a W-shaped profile reflecting edge detection and a U-shaped profile reflecting region filling (Figure 1A), based on previous results (Poort et al., 2012). Consistent with previous analyzes of this dataset, we present average results for two monkeys, while individual results are available in the supplementary materials.

**Figure 1:**
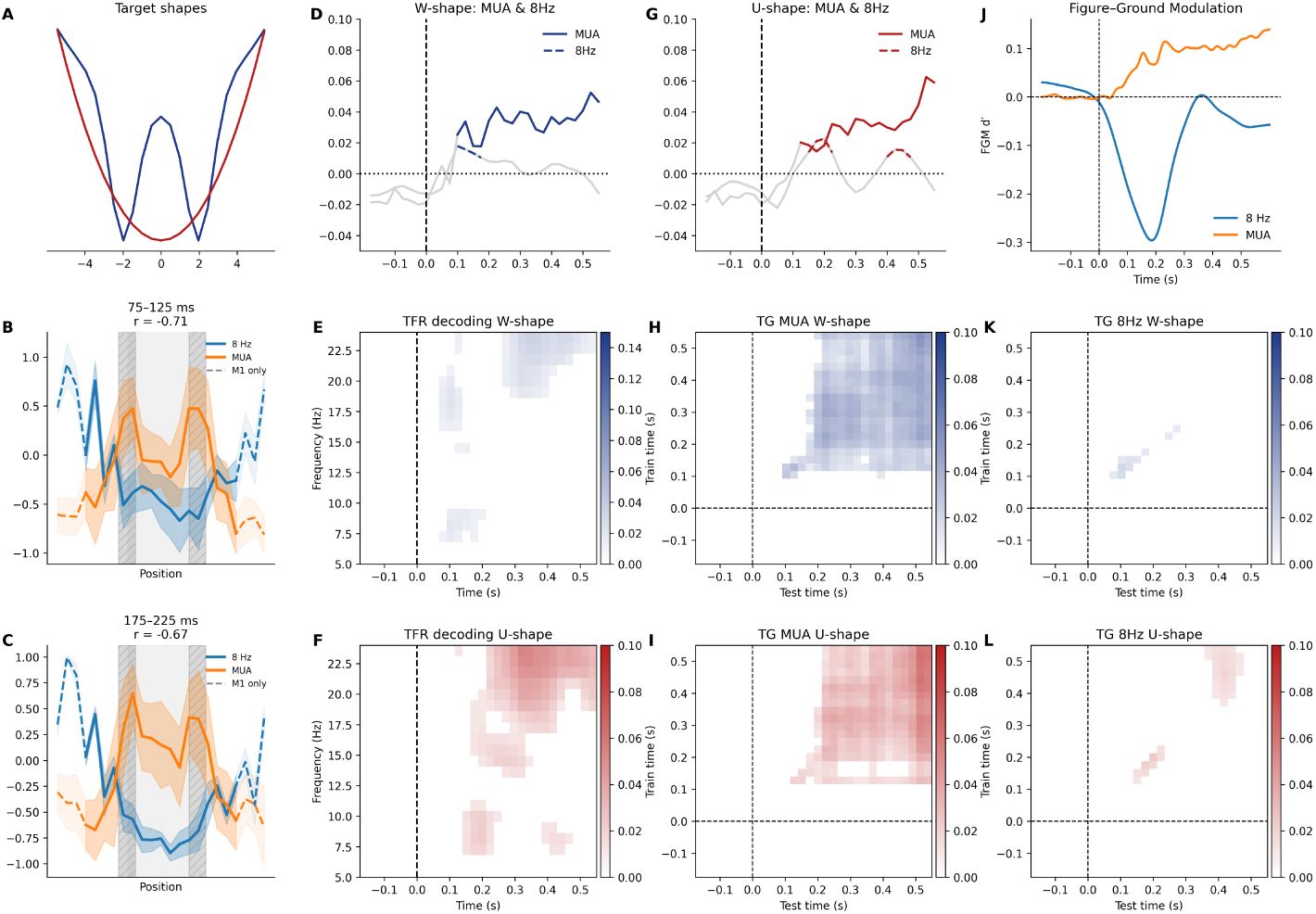
(A) Two testing conditions: blue represent W-shape reflecting edge and center modulation; red represent U-shape reflecting figure filling. (B, C) Averaged MUA and 8 Hz by positions at time windows 100-125 and 175-200 ms (D, G) Decoding results for MUA and LFP 8 Hz for W-shape (blue) and U-shape (red). Solid line MUA, dashed line 8 Hz. (E, F) Decoding results for LFP 5-25 Hz for W-shape (blue) and U-shape (red), colored areas are significant. (H, I) Time generalization of decoding for MUA for W-shape (blue) and U-shape (red), colored areas are significant. (J) Figure-ground modulation orange line MUA, blue line amplitude 8 Hz (K, L) Time generalization of decoding for 8 Hz for W-shape (blue) and U-shape (red), colored areas are significant.

The MUA reliably decoded the position of the figure, with significant decoding emerging approximately 100–125 ms after stimulus onset (Figure 1D,G). Early decoding was stronger for the W-shape than the U-shape and emerged earlier, consistent with the finding that edge-related modulation dominates within the 100–200 ms window. Both profiles peaked at 525 ms, with the U-shape reaching a slightly higher peak (W-shape: *R*^2^ = 0.05 *p <* 0.001; U-shape: *R*^2^ = 0.06, *p <* 0.001), consistent with an increase in figure–ground modulation before saccade onset.

LFP activity in the low frequencies (5–25 Hz) also reliably encoded figure position (Figure 1E, F). For the U-shape, strongest decoding occurred at 25 Hz around 325 ms (*R*^2^ = 0.05, *p <* 0.01); for the W-shape it peaked at 25 Hz around 350 ms (*R*^2^ = 0.03, *p <* 0.01). Within the alpha-theta band (5–15 Hz), both shapes decoded best at 8 Hz but with distinct temporal profiles: W-shape decoding spanned 100–175 ms (peak at 100 ms, *R*^2^ = 0.02, *p <* 0.01), whereas U-shape decoding occurred in two intervals, 150–225 ms and 400–475 ms (peak at 200 ms, *R*^2^ = 0.02, *p <* 0.01). This transient dissociation coincides with the window in which edge- and figure-related modulation are most distinct.

### Time-generalization

To characterize how identified neural representations relate to distinct stages of figure–ground modulation and to assess the temporal stability of alpha-band activity, we performed temporal generalization analysis (King & Dehaene, 2014). A decoder trained at each time point was tested on all other time points, with sustained off-diagonal generalization indicating a stable underlying representation and restricted generalization indicating a transient one.

MUA decoding showed sustained cross-temporal generalization emerging after approximately 225 ms for both shapes (Figure 1H,I). Early responses around 100 ms did not generalize to later time points for the W-shape, and only partially generalized for the U-shape within 100–200 ms. At 8 Hz, generalization was significant but temporally restricted: W-shape between 100–150 ms, U-shape between 200–225 ms (Figure 1K,L). This suggests a transient role for alpha activity during the transition from early edge processing to stable figure representations.

### Position-level correlations

Previous analyzes of this dataset also demonstrated a general increase in alpha power (5-15 Hz) in the OUT position compared to the IN position and the opposite effect for MUA (Poort et al., 2012; van Kerkoerle et al., 2014). Here, we examined this relationship at a finer spatial and temporal scale, focusing on two time windows based on our decoding: an early window (100-125 ms) corresponding to the W-shape peak at 8 Hz, and a later window (175-200 ms) corresponding to the U-shape peak.

We tested the correlation between MUA and previously obtained frequencies of interest averaged across positions. In the early window, 8 Hz amplitude and MUA were strongly negatively correlated across positions (*r* = *−*0.71, *p <* 0.01), in the later window (*r* = *−*0.67, *p <* 0.01) consistent with our prediction and interpretation of alpha as inhibitory signal (Figure 1B,C).

Next, following our previous analysis of this dataset, we calculated figure–ground modulation (FGM) for frequencies of interest. The 8-Hz FGM peaked at d’ = 0.30 at 190 ms, exceeding MUA modulation in magnitude but with a narrower temporal profile (Figure 1B).

### Orientation decoding

Following our prediction that low-frequency activity might also carry feature-specific information, we attempted to decode stimulus orientation using the same temporal windows as those used for spatial encoding. To isolate orientation-related signals from figure–ground effects, trials were separated according to whether the figure was positioned inside the receptive field (IN condition) or whether the background occupied the receptive field (OUT condition), the results of which are presented in the supplementary materials.

IN the IN condition MUA decoding peaked at 75 ms (*accuracy* = 0.85) and remained significant across all subsequent windows, consistently exceeding 75% (Figure 2A). LFP showed an early peak at 50 ms (*accuracy* = 0.67) at 25 Hz with weights aligned with MUA, likely reflecting an early evoked visual response rather than feature-specific coding. Focusing instead on the 100–400 ms window, decoding accuracy peaked at 8 Hz at 100 ms (*accuracy* = 0.65), with weights oriented opposite to those of the MUA (Figure 2B), consistent with a suppressive relationship between alpha amplitude and spiking activity.

**Figure 2:**
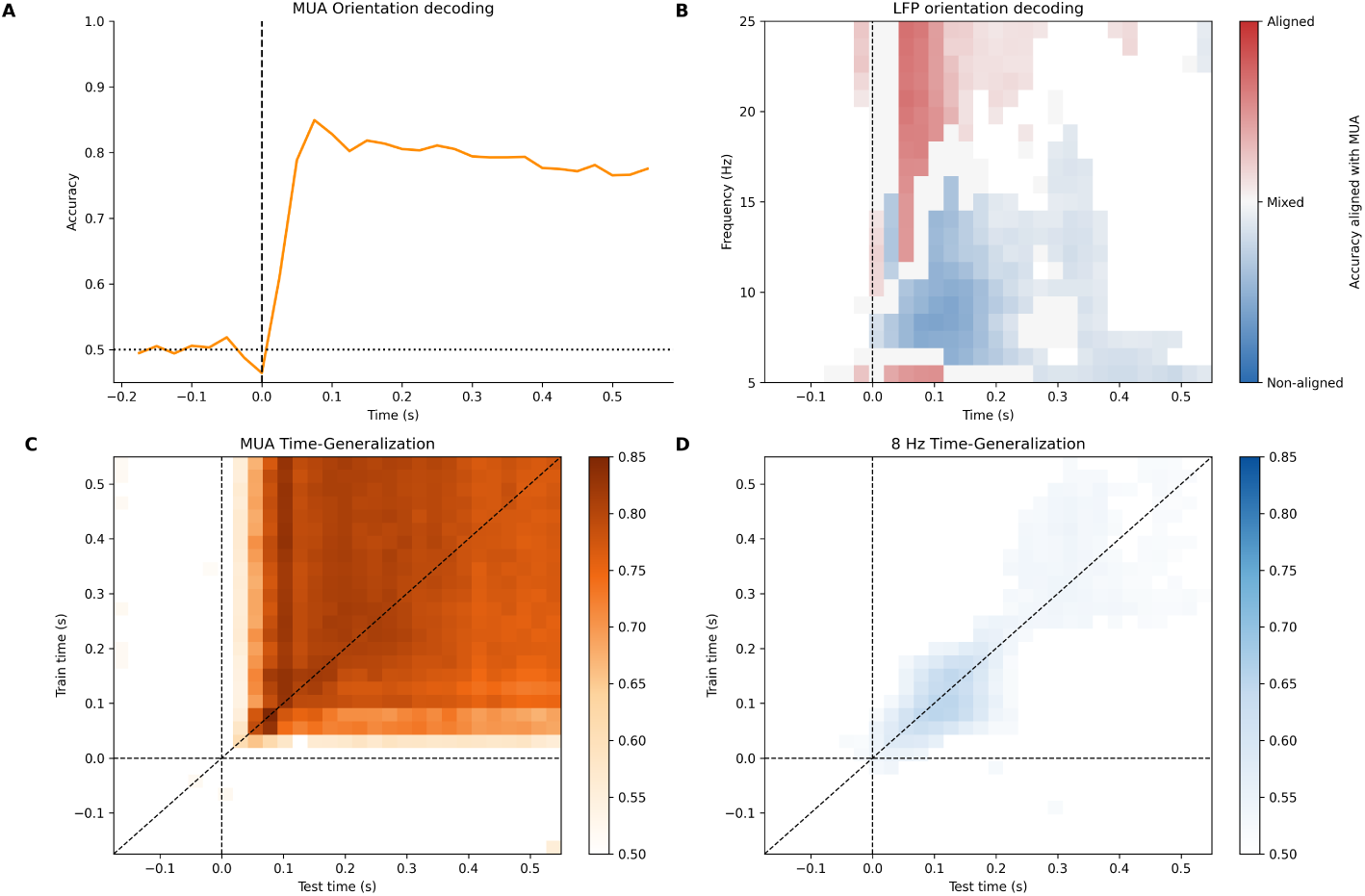
(A) MUA orientation decoding in IN position (B) LFP orientation decoding in IN position, gray area represent significant decoding without stable matching with MUA across channels, blue areas indicate pattern opposite to MUA, and red areas indicate pattern similar to MUA. Color intensity represent normalized accuracy, started from 0.5 to 0.75. (C) MUA orientation decoding time-generalization, colored areas are significant. (D) 8hz orientation decoding time-generalization, colored areas are significant.

### Time-generalization orientation decoding

Repeating our analysis to the position decoding, we carried out time-generalization analysis to check temporal stability of orientation decoding. At IN positions, MUA generalization was sustained and stable after approximately 100 ms with slight decay over time (Figure 2C). At 8 Hz, activity peaking at 100 ms generalized across 25–225 ms but not to later periods; conversely, later activity (250 ms and beyond) did not generalize back to early windows (Figure 2D).

### Amplitude and phase dependence of MUA in V1

The anticorrelation between MUA and 8-Hz amplitude observed during position averaging may indicate both modulation and an independent influence of figure position on alpha activity and MUA amplitude. To expand our analysis and limit the overall distance effect, we examined the dependence of MUA on 8-Hz amplitude and phase separately for the IN and OUT conditions. We conducted analyses in the window of interest (100-200 ms) and in a control window identified for 25-Hz (325-425 ms); results for OUT condition and 25-Hz provided in the supplementary materials.

In the IN condition, MUA was significantly higher in low-amplitude 8 Hz trials than in high-amplitude trials within the 100–200 ms window (Δ = *−*1.67*×* 10*^−^*^6^; *p* = 0.002), with no significant difference in the later window (Δ = *−*5.17*×* 10*^−^*^7^; *p* = 1.0) (Figure 3B).

**Figure 3:**
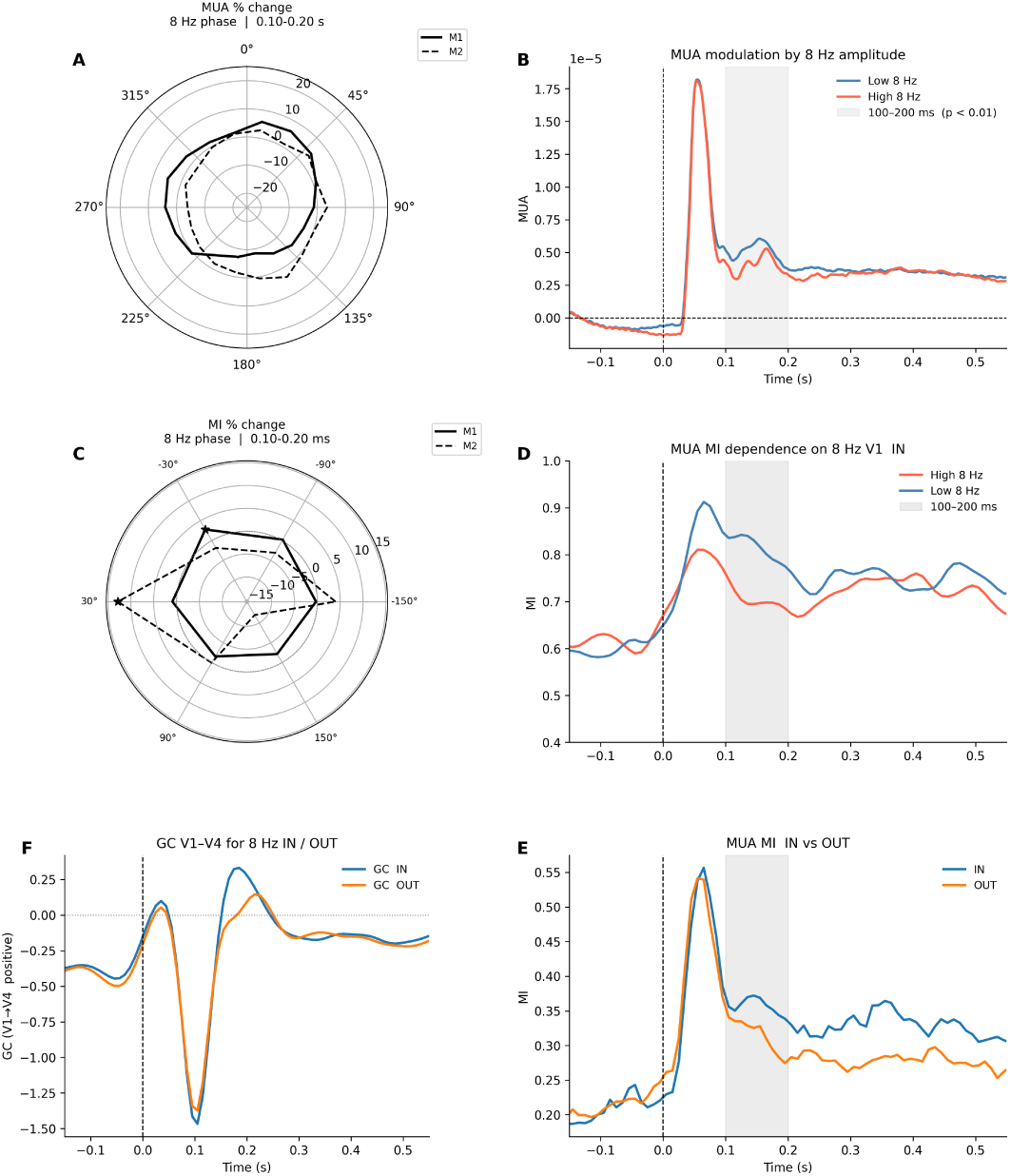
(A) MUA at different phases 8 Hz in the 100 - 200 ms window (B) Blue line MUA at 25% low amplitude 8 Hz trails, orange line MUA at 25% high amplitude 8 Hz trails, gray area represent significant window (C) MI of MUA depending on the phase difference V1-V4 8 Hz, values as a percentage of the average. Solid line represent M1, dash line M2. (D) MI of MUA in In condition orange line MI in low alpha trials, black line MI in high alpha trials. Grey area represent significant window of interest (F) GC direction in time for 8 Hz V1-V4 dominance (positive) V4-V1 dominance (negative). (E) MI of MUA in IN condition (blue line) and OUT condition (orange line). Grey area represent significant window of interest.

We then tested whether MUA was modulated by the phase of 8 Hz oscillations using the modulation index (Tort et al., 2010). Significant phase-dependent modulation was observed in the early window (100–200 ms; M1: *MoI* = 0.002, *p <* 0.001) (Figure 3A) and persisted in the later window (325–425 ms; M1: *MoI* = 0.002, *p <* 0.001).

### Inter-areal interaction between V1 and V4

After establishing the relationship between MUA and 8 Hz amplitude and phase in V1, we next examined whether this modulation affects inter-areal communication between V1 and V4. We quantified coupling strength using Gaussian mutual information (MI) and assessed directionality using Granger causality (GC), focusing on the 8 Hz band in the IN and OUT conditions.

As a validation of our coupling measure, we first compared MI MUA for both conditions. MI was significantly higher in the IN condition than in the OUT during the 100–200 ms window (Δ*MI* = 0.18; *p* = 0.001), indicating stronger inter-areal communication when the figure falls within the receptive field (Figure 3E).

To test whether coupling strength depends on alpha amplitude, trials were divided into high- and low-amplitude groups based on the upper and lower 25th percentiles of 8 Hz amplitude in V1 and then in V4. This approach repeat the analysis used for MUA, MI between low and high trials was compared using permutation testing. Results for the OUT condition are provided in the supplementary materials.

MI was significantly higher in low-amplitude trials in the IN condition (Δ*MI* = *−*0.12; *p* = 0.001) (Figure 3D), consistent with a suppressive role of alpha in regulating inter-areal communication. Repeating the analysis using V4 alpha amplitude yielded the opposite pattern (Δ*MI* = 0.05; *p* = 0.01).

Next, we examined whether MI depends on the instantaneous phase difference between V1 and V4 at 8 Hz. The phase difference was computed trial-by-trial for all V1–V4 channel pairs, summarized as a circular mean across the analysis window, and assigned to six equally-spaced phase bins (60 width). The preferred phase was defined as the bin with the maximum MI averaged across channel pairs, and significance was assessed using a max-statistic permutation test that controls for multiple comparisons across bins.

Significant phase-dependent modulation was observed in both monkeys, though the preferred phase differed between animals and, therefore, results are reported individually. In M1, coupling was strongest in the [*−*60, 0) bin (*MI* = 2.10; *z* = 11.46; *p <* 0.001). In M2, the preferred phase fell in the [0, 60) bin (*MI* = 0.41; *z* = 7.90; *p <* 0.001) (Figure 3C).

### Directionality of alpha-band coupling

To assess the directionality of inter-areal alpha-band communication, we computed Transfer-Reduced Granger Causality (TRGC). Analysis in the 100–200 ms window revealed a marked dominance of the V4*→*V1 direction over V1*→*V4 from approximately 75 to 150 ms after stimulus onset, consistent with top-down feedback modulation of V1 alpha activity (Figure 3F).

### Orientation-selective channels

Finally, to test our hypothesis that alpha activity is stronger in orientation-encoding channels, we identified channels with clear orientation preferences (selective channels) in both monkeys and computed the power difference between preferred and non-preferred orientations across 50-ms windows and all frequencies, using permutation testing with FDR correction.

In the IN condition, the strongest positive difference emerged at 6 Hz around 100 ms, indicating higher power for the preferred orientation and aligning with the MUA results. In contrast, the strongest negative difference—corresponding to lower power for the preferred orientation—was observed at 13 Hz around 150 ms and reached significance within the 100–225-ms interval (Figure 4 B).

**Figure 4:**
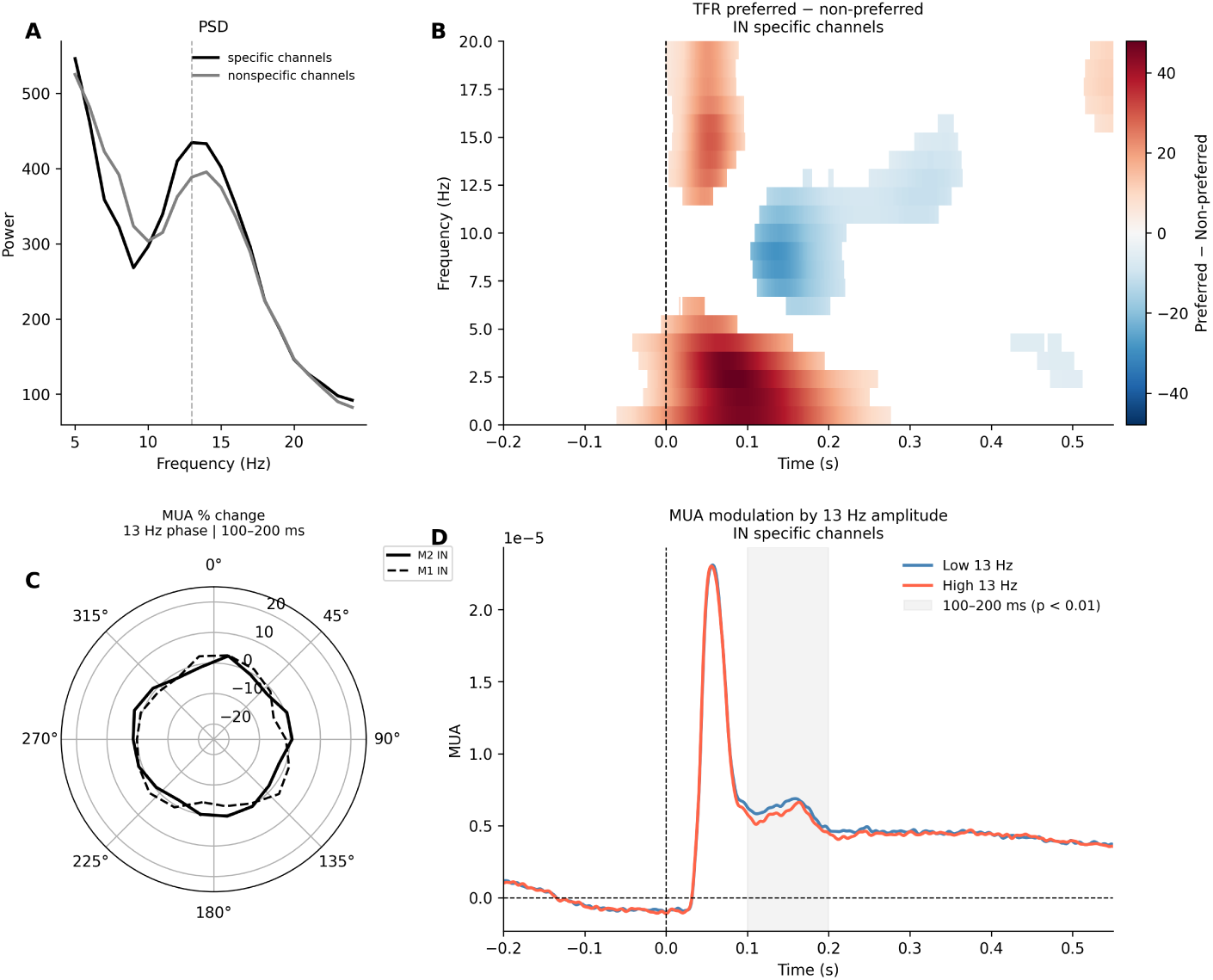
(A) Power spectral density for channels with a preferred orientation (blue solidline) and channels without a preferred orientation (blue dashed line) (B) The difference between the preferred and non-preferred orientations for frequency amplitudes from 5 to 25. Significant windows are marked in color, insignificant ones in white. (C) MUA phasse modulation by 13 Hz (D) MUA amplitude modulation by 13 Hz, gray area represent significant window of interest

Notably, this for selective channels differed from the 8 Hz peak previously identified for orientation decoding across all channels. We therefore compared amplitudes between selective and non-selective channels within the 100–200-ms window. In the IN condition, selective channels exhibited significantly higher amplitudes (Δ = 87; *p <* 0.001) (Figure 4A). We hypothesized that this effect would be attenuated in the OUT condition, given its proposed association with active figure filling-in. Consistent with this prediction, the negative power difference between preferred and non-preferred orientations again peaked at 13 Hz around 150 ms in the OUT condition and remained significant across the same temporal window (Supplementary Figure 10 A). Although the amplitude difference between selective and non-selective channels also remained significant in the OUT condition (Δ = 54; *p <* 0.001) (Supplementary Figure 10 B), the effect size was reduced by 38% relative to the IN condition, suggesting weaker engagement of this frequency band outside the figure representation.

### Amplitude and phase modulation in orientation-selective channels

We next examined whether the 13 Hz amplitude and phase modulate spiking activity and inter-areal coupling in the same manner as the previously identified 8 Hz effect, restricting analyzes to channels with strong orientation selectivity (Figure 4C). In the IN condition, MUA was significantly higher during low-13 Hz-amplitude trials (Δ = 4.73 *×* 10*^−^*^7^; *p* = 0.001) (Figure 4D), consistent with the suppressive relationship between oscillatory amplitude and MUA observed at 8 Hz. In contrast, phase–MUA coupling at 13 Hz was not significant (M1: *MoI* = 0.0002, *p* = 0.60), indicating that 13 Hz activity does not phase-lock spiking within this temporal window.

To test whether 13 Hz amplitude modulated inter-areal coupling, we divided trials into high- and low-amplitude groups and compared MI values using permutation testing. In the IN condition, MI did not differ significantly between amplitude groups (Δ*MI* = *−*0.008; *p* = 0.13). Likewise, no significant phase-dependent modulation of inter-areal MI was observed in either monkey or orientation group. In M1, neither orientation group showed significant effects (OR1: *MI* = 1.97, *z* = *−*1.37, *p* = 0.92; OR2: *MI* = 1.95, *z* = 1.00, *p* = 0.30). Similar null results were observed in M2 (OR1: *MI* = 0.41, *z* = 1.56, *p* = 0.27; OR2: *MI* = 0.41, *z* = 0.67, *p* = 0.89).

## 3 Discussion

Taken together, our results provide evidence in support of the nested oscillation framework proposed by Bonnefond et al. (2017), in which low-frequency activity shapes dynamic information routing through coordinated amplitude modulation and phase synchronization. By combining fine-scale spatial decoding, temporal generalization, amplitude–spike relationships, and inter-areal connectivity analyses, we show that alpha-band activity in V1 carries both spatial and feature-specific information during figure–ground segregation. Moreover, this activity is closely associated with local neuronal excitability within a transient post-stimulus window and with feedback interactions between V4 and V1.

Consistent with the first prediction of the framework, low-frequency activity had fine-scale spatial specificity: figure position could be decoded from alpha amplitude within the 100–225 ms interval after stimulus onset, coinciding with the emergence of figure-ground modulation. These results extend existing evidence for the role of alpha in spatial selectivity and spatial attention (Kelly et al., 2006; Samaha et al., 2016), demonstrating a similar mechanism operating at a finer spatial scale and in the post-stimulus interval rather than solely in anticipatory attention conditions.

The temporal profiles of the two spatial decoding patterns suggest that alpha contributes differently across stages of figure–ground processing. Decoding associated with the edge-dominated W-shaped profile peaked around 100 ms, whereas the figure-filling U-shaped profile peaked later around 200 ms, consistent with a transition from boundary processing to figure representation. The early alpha effect coincided with dominant V4*→*V1 granger causality between 75 and 150 ms, supporting the interpretation of alpha as a feedback-related signal from higher visual areas (Seymour et al., 2019; van Kerkoerle et al., 2014). In particular, this early modulation may reflect feedback from boundary-selective populations in V4, as proposed by Jeurissen et al. (2024), which subsequently contributes to stable figure representations in V1.

Alpha decoding remained temporally transient and did not show sustained temporal generalization, unlike MUA representations, which stabilized after approximately 200 ms. Together with the narrow temporal profile of alpha figure–ground modulation, this shows that alpha primarily supports an early routing or selection process rather than the long-term maintenance of the representation itself. One possible explanation is that alpha-mediated feedback becomes less stable across trials after the perceptual decision and prior to saccade movement, the absence of reaction-time measurements in this dataset prevents direct testing of this possibility. Interestingly, later decoding shifted toward activity around 25 Hz, which may reflect post-decisional or processes associated with frontoparietal feedback and saccade planning (Di Dona & Ronconi, 2023; Engel & Fries, 2010; Fiebelkorn et al., 2018).

The second prediction of the framework — that low-frequency activity modulates local spiking and routes inter-areal communication was also supported by several converging results.

Alpha amplitude was negatively related to MUA across figure positions in the window of interest, and in both IN and OUT conditions low-alpha trials were associated with stronger spiking activity. Similar amplitude–spike relationships were observed for beta-band activity, albeit with later temporal dynamics, suggesting partially overlapping inhibitory mechanisms with distinct functional roles. Together, these results support the interpretation of alpha and beta as inhibitory signals modulating local neuronal output (Jensen & Mazaheri, 2010; Klimesch, 2012) in V1. MUA was additionally modulated by alpha phase in both the early and post-decisional windows, consistent with the idea that low-frequency oscillations organize alternating windows of enhanced and suppressed excitability (Bollimunta et al., 2011; Dougherty et al., 2017; Spaak et al., 2012). We note that the alpha-related effects observed here emerge within 100–200 ms of stimulus onset, temporally close to the initial evoked response at approximately 60–70 ms. Some amplitude- and phase-based measures may therefore partially reflect residual evoked activity. For amplitude, peak decoding latency, frequency specificity, and temporal generalization provide indirect support for a genuine oscillatory contribution. Phase estimates, however, are inherently noisy in short windows containing fewer than one complete cycle, and the evoked response itself can bias instantaneous phase measurements. Indirect support for genuine phase-dependent modulation comes from our results in the post-decisional window (325–425 ms), which also show significant 8 Hz phase dependence of MUA at a time well separated from the evoked response. Future work using task designs that temporally separate evoked and induced components — for example by testing naive animals or using stimuli that engage later and more sustained figure-filling processes (Huang & Paradiso, 2008) — would allow cleaner characterization of low-frequency phase dynamics in this context. The role of alpha in inter-areal communication was further supported by MI results. Inter-areal MI between V1 and V4 MUA began to dissociate around 100 ms and was significantly higher in the IN condition than the OUT condition between 100 and 200 ms, indicating stronger coupling when the figure falls within the receptive field. Within the same window, MI was higher in low-alpha trials in V1 for both conditions, and inter-areal coupling was significantly modulated by the instantaneous V1–V4 phase difference, consistent with a role for alpha synchronization in gating effective communication between areas.

In V4, however, the amplitude–coupling relationship was asymmetric: MI was higher in low-alpha trials in the OUT condition but higher in high-alpha trials in the IN condition. This asymmetry may reflect different functional roles of alpha at different levels of the visual hierarchy. In V1, alpha may primarily reflect inhibitory gating of incoming feedback signals, while in V4 — given its larger receptive fields and stronger top-down drive — alpha may be more closely related to the generation or coordination of feedback. This interpretation is consistent with the observation that V4 alpha behaves similarly across IN and OUT conditions when assessed from V1’s perspective. An alternative account holds that during IN trials, when the figure falls within the V4 receptive field, alpha in V4 selectively suppresses responses to task-irrelevant features such as the background grating, sharpening the figure representation through selective inhibition rather than global suppression. Under this view, higher V4 alpha during IN trials would reflect fine-tuned routing rather than reduced communication. However, V4 alpha decoding was significant only after 150 ms and reached a low peak accuracy of 0.56, which argues against a strong feature-selective interpretation and leaves the mechanistic account of the V4 asymmetry an open question for future work.

Consistent with our third prediction, alpha activity also encoded stimulus orientation within the same temporal window as position decoding, suggesting that alpha supports both spatial and feature-specific representations within a shared temporal regime. For the majority of channels, the peak decoding frequency (8 Hz) was the same as for position decoding. Comparison of decoding weights revealed an opposite pattern across channels relative to MUA, further supporting the nagative correlation between MUA and alpha across positions and consistent with the high- and low-alpha trial results.

In line with Clausner et al. (2025), we confirmed that orientation-selective channels showed a peak difference between preferred and non-preferred orientations in high alpha (13 Hz), and also demonstrated that these channels exhibited higher 13 Hz amplitude than non-selective channels. This may reflect an increased sampling rate for task-relevant features at higher alpha frequencies. Although higher alpha amplitude might appear to contradict the suppressive role of alpha described above, these results are not mutually exclusive: higher amplitude in orientation-selective channels may reflect fine-tuned inhibitory suppression of the non-preferred orientation, which is consistent with the finding that low-alpha trials showed significantly higher MUA. Phase modulation at 13 Hz was not significant, and inter-areal MI results were also null at this frequency. However, these null results should be interpreted with caution, as the Gaussian MI estimator relies on cross-covariance matrices and is sensitive to the ratio of observations to channels; future studies with larger channel sets and a broader range of orientation features would provide a more sensitive test.

Although our results suggest a role for alpha in fine-scale information routing, our findings are based on data where different RF positions correspond to different individual trials, rather than within a single recording. Such an analysis is limited by amplitude variability across trials and may underestimate the fine-grained regulation of individual trials. The recording design assumed a stable receptive field distance for V1 and V4 channels, enabling comparison of IN and OUT conditions for both areas simultaneously, but not intermediate configurations. Additionally, the U-shaped decoding profile, which reflects figure filling, assumes that alpha modulation increases with distance from the RF center. Although this was supported by our position-averaging results, we hypothesize that alpha modulation may be local and could plateau or decrease at greater distances, a question that would require recordings spanning a larger range of receptive field positions.

Finally, although the results are presented as averages across monkeys and individual results broadly agree with the group-level findings, some discrepancies are worth noting. The peak alpha frequency for position decoding differed slightly between animals (M1: 9 Hz; M2: 10 Hz), and significant 25 Hz decoding was observed only in M1, though cross-position correlations were significant in both monkeys. The amplitude dependence of MUA was absent or weak in M2 which, together with the smaller FGM peak, may reflect differences in attentional strategy or general individual differences in observations.

## 4 Methods

In this study, we reanalyze a previously published dataset containing local field potentials (LFP) and multi-unit activity (MUA) recordings from visual areas V1 and V4 in two macaque monkeys (Poort et al., 2012).

The monkeys in this dataset performed the figure-ground segregation task. Each trial began with the monkey fixing a central red point on a gray background. After 300 ms of stable fixation, the visual stimulus appeared. The stimulus consisted of three elements: (i) a background texture, (ii) a 4° × 4° square figure presented in the lower visual field, and (iii) two curved line elements in the upper visual field used for the additional control condition (Figure 5A). Both the figure and background were composed of oriented line segments (45° and 135°), presented in random order.

**Figure 5:**
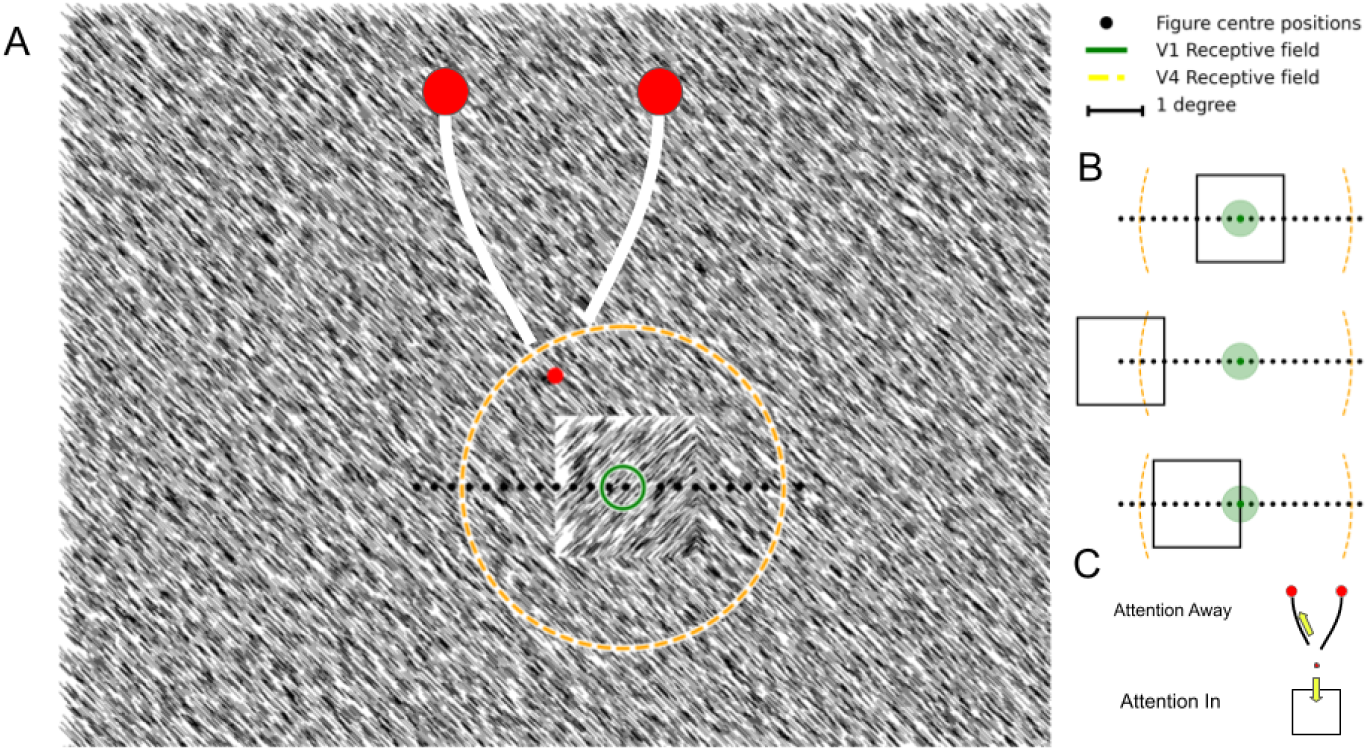
(A). Texture segregation stimulus with an orientation-defined square figure superimposed on a background with an orthogonal orientation, and two curves connected to larger red circles. The smaller red circle in the center is the fixation point. The green circle is the RF of a V1 recording site, the orange circle is the 50% RF of a V4 recording site. The black dots are the positions of the figure center, which have been used in the experiment. (B) The square Figure (4° × 4°) could appear at one of 23 locations for M1 and 17 locations for M2. Black dots represent possible locations of the figure center. In some of the conditions the RF V1 (green circle) fell on the center (upper), on the background (middle) or on the edge (bottom). The orange circle shows the boundaries of 50% RF V4. (C) Attention In and Attention Away conditions. Attention In condition required a saccade to the center of the square figure. Attention Away condition required a saccade to the circle that was connected to the fixation point by one of the curves.

Monkey was required to make a saccade into a 2.5° × 2.5° target window centered on the middle of the figure. The position of the figure was systematically varied along the spatial horizontal X relative to the center of the receptive field (RF) of the recorded sites in steps of 0.5°. In total, 23 positions were tested for monkey 1 and 17 for monkey 2 (RF center relative to figure center ranging from –5.5° to +5.5°). All figure positions were presented in a randomly inter-leaved sequence in both tasks. Trials in which the figure fell into the receptive field (RF) were classified the IN condition. Trials in which both the figure and the background fell into the receptive field were defined as the Edge condition. Finally, trials in which the figure was outside the receptive field were designated as the Out condition. Correct responses in both tasks were rewarded with apple juice. On each experimental day, the monkey performed only one of the two tasks, the task was cued at the start of the session. Performance accuracy was high: M1: 97% (figure detection), 92% (curve tracing) M2 J: 99% and 96% respectively.

### Preprocessing

Electrophysiological data (LFP and MUA) were preprocessed in MATLAB (R2021a, MathWorks) following the pipeline described in van Kerkoerle et al. (2014). Only valid trials were retained for analysis. Channels were included if their stimulus-evoked response exceeded 3 standard deviations above pre-stimulus baseline activity. The noise from the power line was removed using notch filters at 50, 100 and 150 Hz. For each trial, baseline was corrected by subtracting the mean signal before stimulation from *−*200 to *−*50 ms.

### Time-frequency analysis

LFP time–frequency analysis was performed in the FieldTrip toolbox using the superlet transform. The frequency range of interest was set from 5 to 25 Hz (cfg.foi), covering the theta, alpha, and beta frequency bands. The base wavelet width was set to 3 cycles (cfg.width), providing a balance between temporal and spectral resolution. Wavelet length was determined using a Gaussian envelope with a width parameter of 3 (cfg.gwidth), reducing spectral leakage between neighboring frequencies. Superlets were combined using the multiplicative approach (cfg.combine = ‘multiplicative’), which provides higher frequency resolution compared to the additive method. Time–frequency representations were computed for the interval from –0.2 to 0.6 s relative to stimulus onset.

### Position decoding analysis

To examine the temporal dynamics of neural representations, we performed a time-resolved decoding analysis using sliding windows of 50 ms duration, advanced in steps of 25 ms, spanning –200 to 600 ms relative to stimulus onset. Within each window, neural activity was averaged across time, yielding trial-wise feature vectors (trials × channels) for each frequency band.

Stimulus identities were mapped onto continuous representational profiles reflecting their positions within a predefined stimulus space derived from previously reported MUA response profiles (Poort et al., 2012). Specifically, stimulus labels were converted into distances within this representational space, and two target functions were defined to capture distinct hypothesized neural geometries: a W-shaped profile associated with early boundary detection and a U-shaped profile associated with later region-filling processes (Figure 1A).

Stimulus identities were mapped to continuous representational profiles reflecting their positions in a predefined stimulus space, derived from previously obtained MUA response profiles (Poort et al., 2012). Specifically, stimulus labels were converted to distances within this space, and two target functions were constructed to capture distinct hypothesized representational geometries: a W-shaped profile associated with early boundary detection and a U-shaped profile associated with late region filling (Figure 1A). The W-shape target profile was defined as:

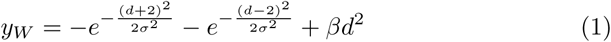

where *d* denotes the stimulus distance, *σ* = 0.7 controls the width of the side components, and *β* = 0.02 determines the contribution of the quadratic term. With these parameters, the modulation associated with edge representations was approximately 40% greater than that of the central representation.

The U-shape target profile was defined as the squared distance function:

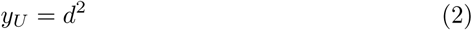

Decoding was performed using ridge regression with leave-one-session-out (LOSO) cross-validation across 3 session for M1 and 5 for M2. To avoid biases related to unequal sampling, trial numbers were equalized across stimulus positions and orientations by random subsampling. In each fold, trials from one recording session were held out for testing, while the model was trained on data from the remaining sessions.

Prediction performance was quantified using the coefficient of determination (R²) between predicted and true target values. Statistical significance was assessed using permutation testing (n = 1000), performed independently for each time window and frequency band. The analysis was conducted separately for the two stimulus orientations, after which observed and permuted R² values were averaged across orientations. P-values were computed as the proportion of permutation scores exceeding the observed R² and were corrected for multiple comparisons using the false discovery rate (FDR). For analyses combining data across monkeys, the mean R² across animals was compared against the averaged permutation distribution.

After identifying the frequencies of interest, we performed a temporal generalization (TG) analysis to investigate the stability and temporal evolution of the decoded representational structures across time. Ridge regression models were trained to predict the representational target values (W-shape or U-shape) using leave-one-session-out (LOSO) cross-validation and the same temporal windowing procedure as in the main decoding analysis. For each pair of training and testing time windows, activity was averaged within the corresponding windows, and the trained model was used to generate predictions for the held-out data. Prediction performance was quantified using the coefficient of determination R² between predicted and true target values, resulting in a train-time × test-time generalization matrix. Statistical significance was assessed using permutation testing (*n* = 1000). Analyses were performed separately for the two stimulus orientations, after which observed and permuted R² values were averaged across orientations. P-values were calculated as the proportion of permutation scores exceeding the observed R² values and were corrected for multiple comparisons using the false discovery rate (FDR).

### Orientation decoding

For orientation decoding, we used MUA data from IN positions. Time windows were identical to those used for position decoding, and the data were balanced across conditions. We applied the same leave-one-session-out (LOSO) cross-validation procedure, but used a linear classifier instead of linear regression.

To find channel with preferred orientation, we computed Haufe-transformed weights within the 0.1–0.2 s time window (Haufe et al., 2014). Decoding performance was quantified using classification accuracy. Statistical significance was evaluated using permutation testing (*n* = 1000*permutations*), performed independently for each time window. Based on their contribution to orientation decoding in IN and OUT conditions, channels were grouped into three categories: rientationselective channels, and non-selective channels.

To compare the spatial organization of orientation-selective representations across signals and frequency bands, we quantified the similarity between MUA decoding weights and LFP decoding weights using cosine similarity. For each time window and frequency band, cosine similarity was computed between the Haufe-transformed MUA weight vector and the corresponding LFP weight vector separately for each monkey. This analysis provided a measure of the extent to which frequency-specific LFP representations aligned with spatial pattern of orientation selectivity observed in MUA activity.

To assess whether orientation-selective channels exhibit higher alpha amplitude than non-selective channels, we compared mean amplitude within the 100–200 ms window of interest between two groups of channels. For each monkey, amplitude was extracted at the frequency of interest by averaging across the time window and selected channels, yielding a single amplitude value per trial. The observed difference in mean amplitude between selective and non-selective channels was computed separately for each monkey and then averaged across monkeys. Statistical significance was assessed using a two-sided permutation test (*n* = 1000). For each permutation, trial labels were randomly reassigned between the selective and non-selective pools within each monkey, the mean difference was recomputed, and the results were averaged across monkeys to form a null distribution.

To characterize the frequency and temporal profile of orientation selectivity across channels, we computed the difference in spectral amplitude between preferred and non-preferred orientations across all time–frequency bins. Differences were averaged across channels and trials separately for each monkey. Statistical significance was assessed using a permutation test (*n* = 1000). For each permutation, the sign of each trial’s difference was randomly flipped, and the mean across trials was recomputed, yielding a null distribution at each time–frequency point. The p-value at each point was defined as the proportion of permuted means whose absolute value equalled or exceeded the observed absolute difference. Multiple comparisons across the time–frequency plane were corrected using the FDR.

### Figure-ground modulation

We also compute figure–ground modulation using the same approach as in previous analyzes of this dataset (Poort et al., 2012). Figure–ground modulation of MUA (MUA-FGM) was computed as the normalized difference between responses to figure and background conditions, defined as the difference in mean responses divided by the pooled standard deviation across the two conditions.

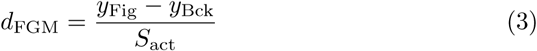

We extended this analysis to low-frequency LFP activity within the frequency bands of interest, where we expected an effect opposite to that observed for MUA. The metric was computed separately for each channel and each orientastion.

### Correlation by positions

For each channel, neural activity was averaged across stimulus positions (n = 23) using the peak frequencies identified by the ridge regression model, as well as MUA. The mean and standard deviation were then computed across channels, and results were averaged across monkeys. Within the time window of interest, we computed the correlation between low-frequency amplitude and MUA across positions, with multiple comparisons corrected using the discovery rate (FDR). As only 17 stimulus positions were common across all sessions for the M2, analyses requiring a consistent position set were restricted to data from a M1 for the remaining six positions.

### Ampitude MUA modulation

We calculated MUA modulation by dividing trials under both IN and OUT conditions separately into 25 % high and 25 % low amplitude trials for the frequency of interest for each window (50 ms width). Then we calculated the difference between the averaged MUA across channels for each time window. For two global windows of interest (100-200 ms and 325-425 ms), we calculated the difference between MUAs between trial groups and compared them with the absolute permutation difference (n = 1000) for these conditions. Multiple comparisons across time windows were then controlled using false discovery rate (FDR) correction.

### Phase MUA coupling

To quantify phase–MUA coupling, we calculated the modulation index (MoI) using the procedure described by (Tort et al., 2010) and adopted in prior studies (e.g., (Dougherty et al., 2017), restricting the analysis to the predefined time window of interest. The oscillatory phase range (*−π, π*) was partitioned into 18 equal bins, and MUA values were averaged within each bin, yielding a phase-dependent amplitude distribution.

The modulation index was defined as the Kullback–Leibler divergence between this empirical phase-conditioned MUA distribution and a uniform distribution, normalized to obtain MoI. MoI values were first computed for each channel individually and then averaged across channels. Statistical significance was evaluated via a permutation test (*n* = 1000), in which MUA samples were randomly reassigned with respect to phase, producing a null distribution of MoI values. Population-level p-values were obtained by comparing the observed MoI to this null distribution.

### Mutual information

We quantified functional coupling between V1 and V4 population activity using a multivariate Gaussian copula mutual information (GCMI) estimator (Ince et al., 2017; Wen et al., 2025). Mutual information (MI) was computed in sliding temporal windows (50 ms width, 25 ms step) under a multivariate Gaussian assumption:

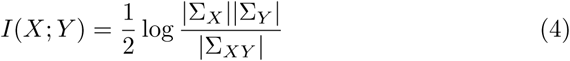

where Σ*_X_*, Σ*_Y_*, and Σ*_XY_* denote the covariance matrices of V1 activity, V4 activity, and their joint concatenation, respectively.

To account for transmission delays between V1 and V4, we introduced a fixed temporal lag of 17 and 20 ms and when computing cross-area covariances. These lags were chosen based on the observed latency differences in visual responses across monkeys.

For frequency-specific analyzes (8 Hz and 25 Hz conditions), we tested three temporal alignments ( 20*/*17*ms*, 0*ms*, + 20*/*17*ms*) to ensure robustness. In the 8 Hz condition, the 20 ms lag produced the highest average mutual information (MI) and was therefore selected for all subsequent analyses.

To examine how inter-areal coupling depends on alpha-band activity, trials were split into high- and low-alpha groups based on the upper and lower 25th percentiles of alpha power in the IN and OUT conditions. MI was computed separately for each group using the same procedure as above. We then focused on the 100–200 ms time window, calculated the group difference, and averaged the results across monkeys. Statistical significance was assessed using a permutation test (*n* = 1000*iterations*). Multiple comparisons were corrected across conditions and areas (V1 and V4).

Granger causality (GC) was estimated using the MNE-Connectivity toolbox in the 5–25 Hz frequency range. To control for potential biases due to temporal asymmetries and spurious directionality, we repeated the analysis on time-reversed signals and subtracted the resulting GC values.

### Phase modulation of the MI

To assess how the V1–V4 phase difference modulates inter-areal coupling, we computed the instantaneous phase difference at 8 Hz for each trial and each V1–V4 channel pair within the 100–200 ms window of interest. Trials were assigned to six equally spaced phase bins (60 width), with the number of trials per bin equalized across bins to prevent bias in MI estimation. Mutual information between V1 and V4 MUA was computed for each bin using all channels simultaneously, and the resulting MI profiles were averaged across channel pairs. The preferred phase was defined as the bin with the maximum mean MI. Statistical significance was assessed using a max-statistic permutation test (*n* = 1000): for each permutation, phase labels were randomly shuffled within each channel pair, MI was recomputed per bin, and the maximum across bins was retained. The null distribution was obtained by averaging these per-pair maxima across channel pairs, and the observed maximum was compared against this distribution to yield a *z*-score and *p*-value.

## Author Contributions

Conceptualization: M.B., D.S.

Methodology: M.B., D.S.

Software: D.S., G.A.

Validation: M.B., D.S.

Formal analysis: D.S., G.A.

Writing – Original Draft: M.B., D.S.

Visualization: D.S.

Supervision: M.B.

Funding acquisition: M.B.

## Data and Code Availability

Code will be made available in an open-access repository upon publication.

## Supplementary Materials

Orientation prediction analysis in V4 revealed a peak decoding performance at 8 Hz (accuracy = 0.56) approximately 100 ms after stimulus onset.

**Figure S1:**
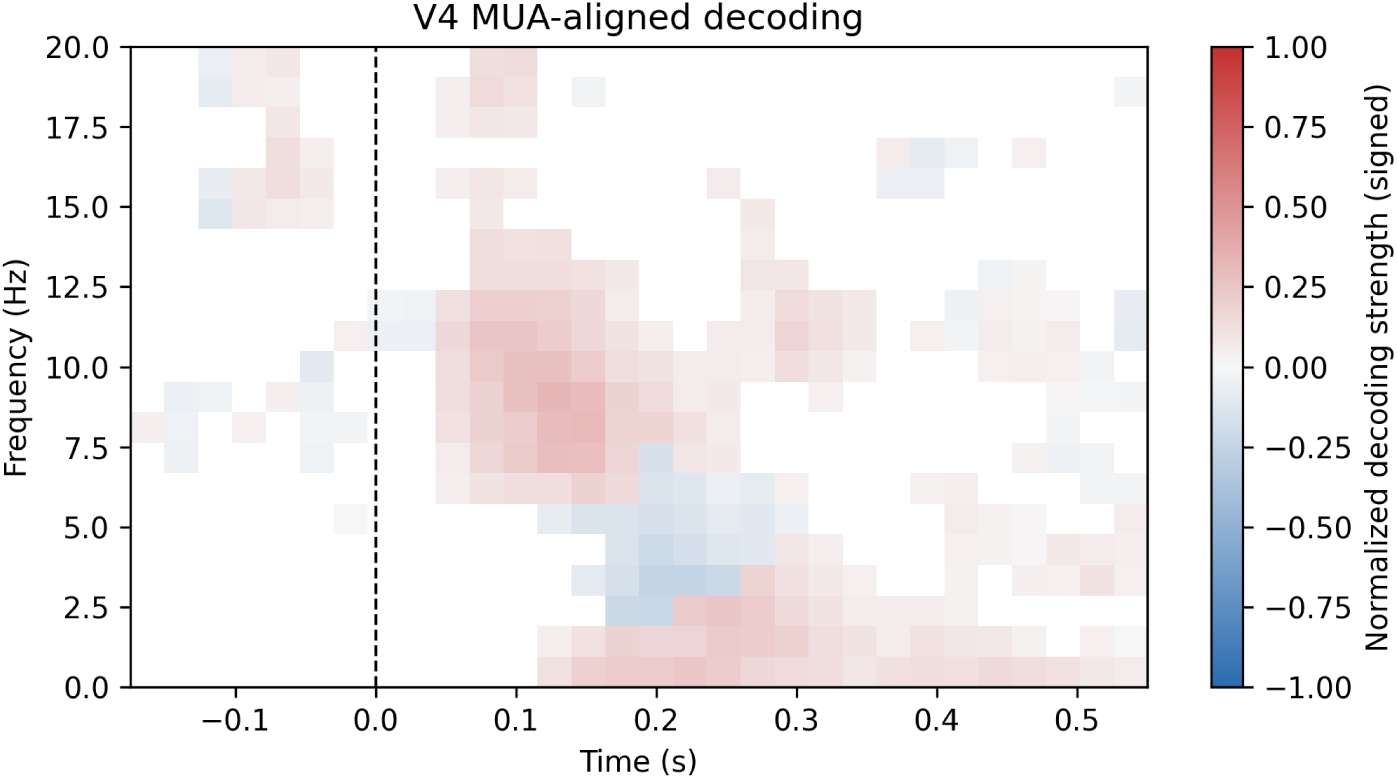
Orientation decoding performance in V4.

Additional analysis of the orientation-specific channels’ mutual information revealed no significant differences between the high and low 13 Hz frequencies.

**Figure S2:**
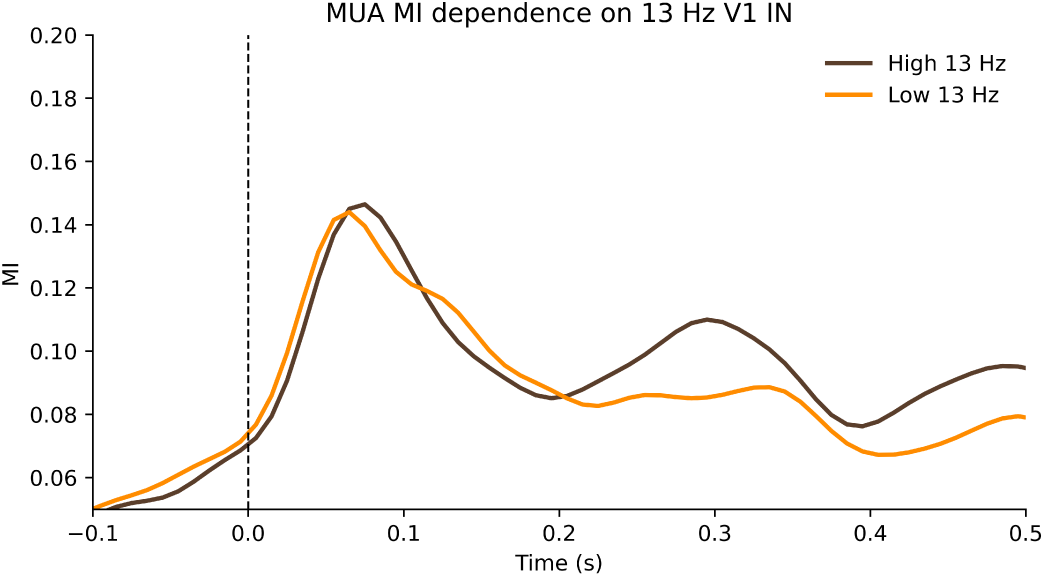
MI of MUA in In condition orange line MI in low alpha trials, black line MI in high alpha trials. Grey area represent significant window of interest.

### S1. Results for the OUT condition

#### Orientation decoding

MUA decoding peaked at 75 ms (*accuracy* = 0.76), mirroring the IN result but with lower sustained accuracy. LFP showed an early peak at 50 ms (*accuracy* = 0.72) at 25 Hz with weights aligned with MUA. Applying the 8 Hz frequency identified at IN positions, decoding peaked at 100 ms (*accuracy* = 0.66), but with mixed weight orientations rather than the clear opposition seen at IN positions. Cross-temporal generalization yielded a closely parallel pattern to the IN condition: MUA generalization was sustained after 100 ms, and 8 Hz activity generalized across 25–225 ms but not to later periods, suggesting a common mechanism underlying 8 Hz orientation coding in both conditions.

**Figure S3:**
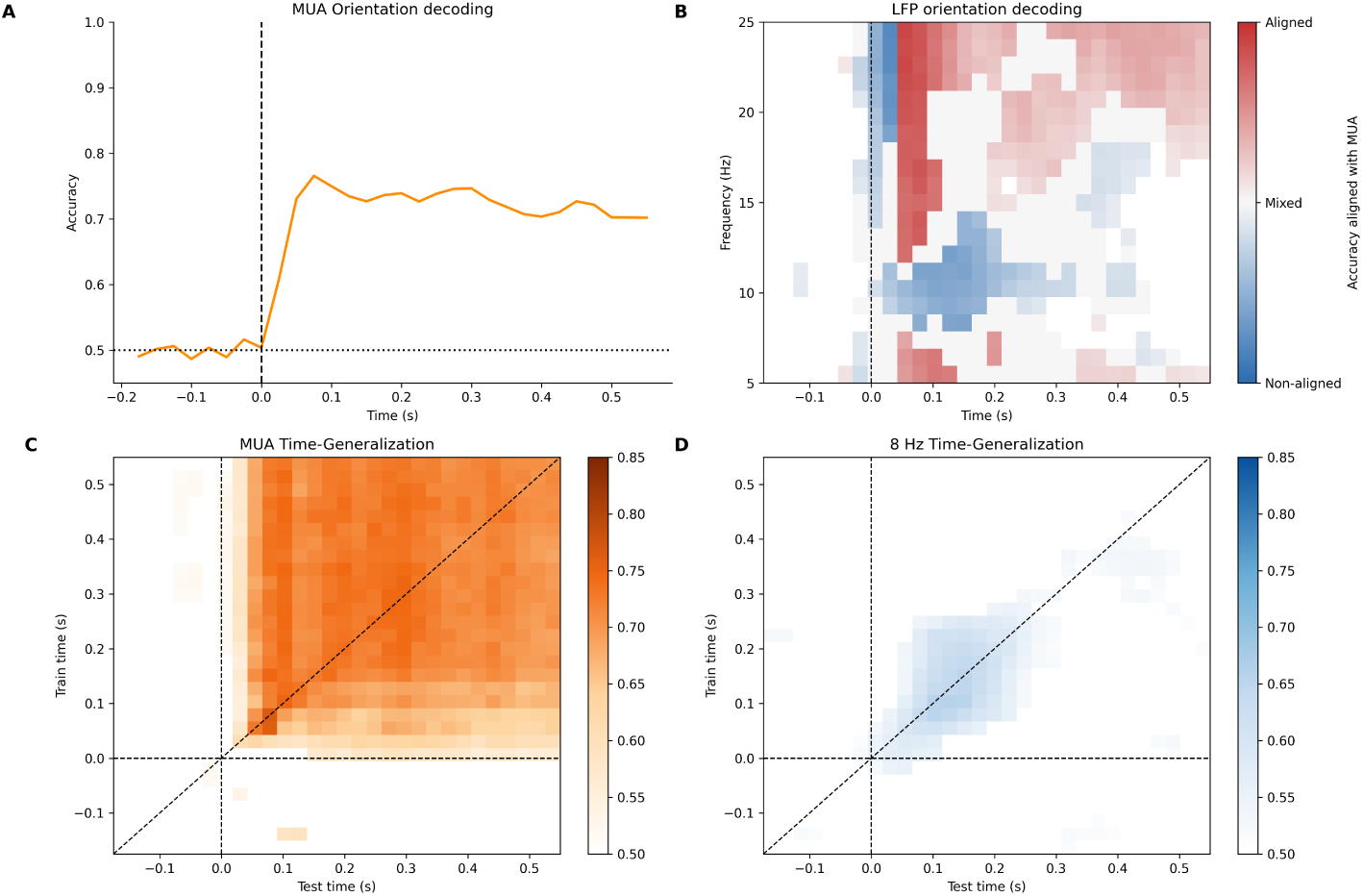
(A) MUA orientation decoding in IN position (B) LFP orientation decoding in IN position, gray area represent significant decoding without stable matching with MUA across channels, blue areas indicate pattern opposite to MUA, and red areas indicate pattern similar to MUA. Color intensity represent normalized accuracy, started from 0.5 to 0.75. (C) MUA orientation decoding time-generalization, colored areas are significant. (D) 8hz orientation decoding time-generalization, colored areas are significant.

#### Amplitude modulation

In the OUT condition, MUA was significantly higher in low-amplitude 8 Hz trials than in high-amplitude trials in the early window (100–200 ms; Δ = 1.82 *×* 10*^−^*^6^, *p* = 0.002), with no significant difference in the later window (Δ = *−*5.17 *×* 10*^−^*^7^, *p* = 1.0).

#### Phase-dependent modulation

At 8 Hz, phase modulation remained robust in the early window (M1: *MoI* = 0.006, *p <* 0.001; M2: *MoI* = 0.002, *p <* 0.001). In the later window, the effect persisted in M2 (*MoI* = 0.010, *p* = 0.002) but not in M1 (*MoI* = 0.001, *p* = 0.89). No significant phase modulation was observed at 25 Hz in either time window (early: M1 *MoI* = 0.0004, *p* = 0.719; M2 *MoI* = 0.0007, *p* = 0.348; late: M1 *MoI* = 0.0018, *p* = 0.573; M2 *MoI* = 0.0025, *p* = 0.530).

**Figure S4:**
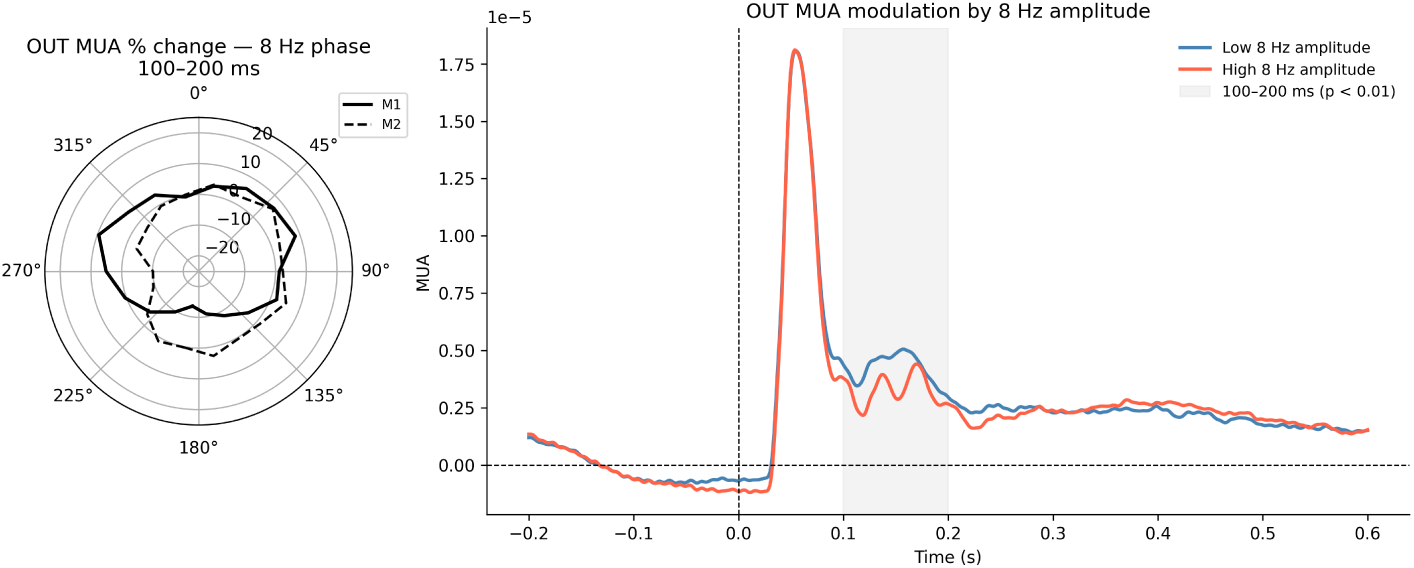
(A) MUA at different phases 8 Hz in the 100 - 200 ms window (B) Blue line MUA at 25% low amplitude 8 Hz trails, orange line MUA at 25% high amplitude 8 Hz trails, gray area represent significant window

#### Inter-areal coupling

MI between V1 and V4 MUA was higher in low-alpha trials in the OUT condition (Δ*MI* = *−*0.11; *p* = 0.001), consistent with a suppressive role of alpha in regulating inter-areal communication. Repeating the analysis using V4 alpha amplitude yielded the same direction for OUT trials (Δ*MI* = *−*0.08; *p* = 0.002). Phase-dependent modulation of inter-areal MI was significant in both monkeys: M1 preferred phase [120, 180) (*MI* = 1.30; *z* = 9.59; *p <* 0.001); M2 preferred phase [0, 60) (*MI* = 0.44; *z* = 9.09; *p <* 0.001).

### Orientation-selective channels

Results regarding specific channels in the OUT condition are provided in the main text.

### S2. Results for 25 Hz

#### Position decoding

The 25 Hz band showed no significant position–MUA correlation in the early window (*r* = *−*0.30, *p* = 0.46), but a significant negative correlation emerged in the later window (*r* = *−*0.60, *p <* 0.01). Figure–ground modulation at 25 Hz peaked later at *d^′^*= *−*0.33 at 425 ms, consistent with its selective emergence in the late time window.

**Figure S5:**
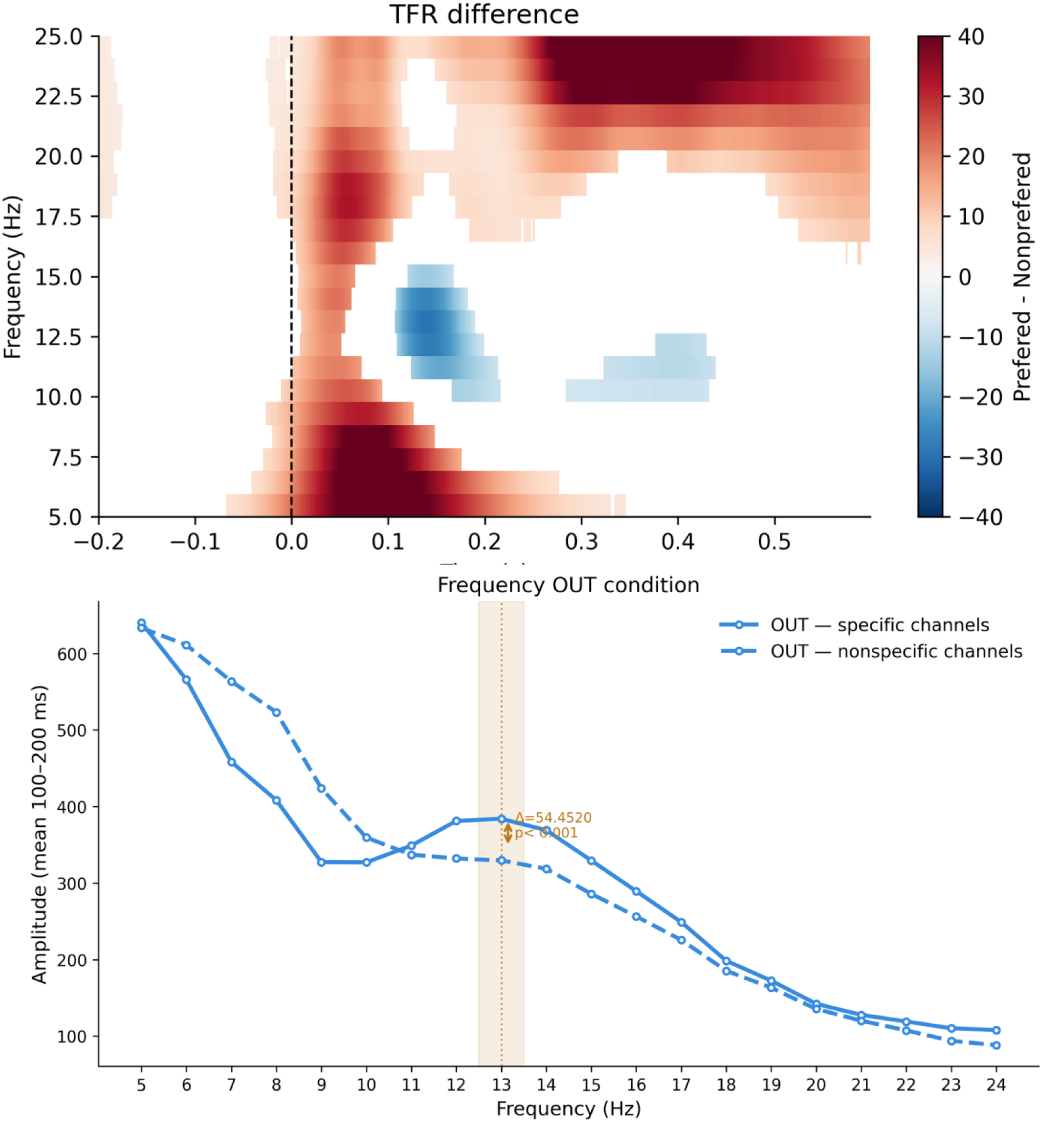
(A) The difference between the preferred and non-preferred orientations for frequency amplitudes from 5 to 25. Significant windows are marked in color, insignificant ones in white. (B) Power spectral density for channels with a preferred orientation (blue solidline) and channels without a preferred orientation (blue dashed line).

#### Amplitude modulation — IN condition

A significant amplitude-dependent MUA difference was present in both the early (Δ = 4.0 *×* 10*^−^*^7^, *p* = 0.007) and later windows (Δ = 4.6 *×* 10*^−^*^7^, *p* = 0.002).

#### Phase-dependent modulation — IN condition

Phase modulation at 25 Hz was inconsistent across monkeys. In the early window, no significant modulation was observed in M1 (*MoI* = 0.0002, *p* = 1.0), whereas M2 showed a significant effect (*MoI* = 0.001, *p* = 0.004). In the later window, neither monkey reached significance after correction (M1: *MoI* = 0.0003, *p* = 1.0; M2: *MoI* = 0.002, *p* = 0.10).

### S3. Individual monkey results

Results are presented separately for each monkey to allow direct comparison with the group-level analyses.

**Figure S6:**
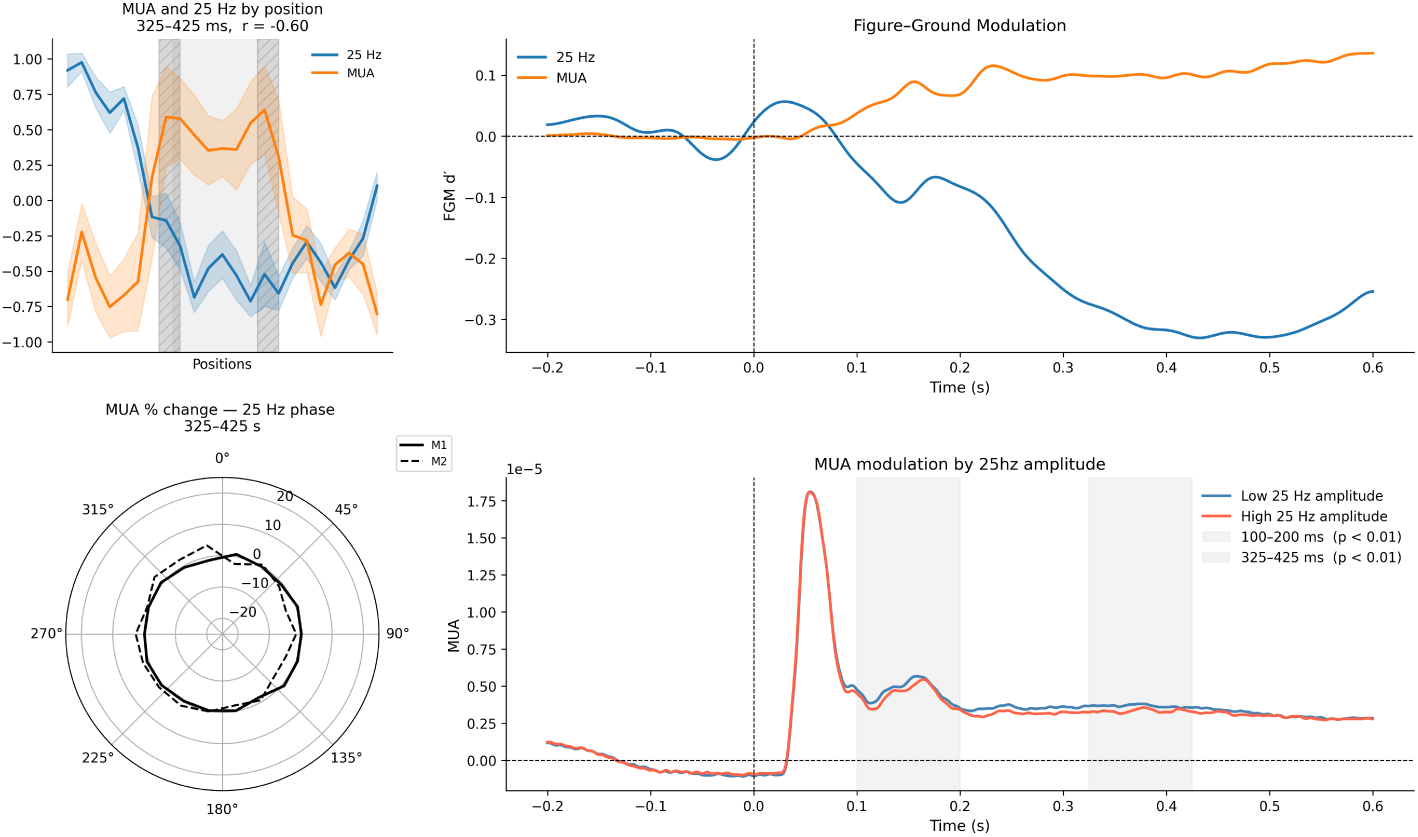
(A) Averaged MUA and 25 Hz by positions at time windows 325-425 ms, Gray area when the figure is in position IN dashed represent edge positions. (B) Figure-ground modulation MUA (orange line) and 25Hz (blue line) (C) MUA phasse modulation by 25 Hz for time window 325-425 ms. Solid line represent M1, dashed line represent M2 (D) MUA amplitude modulation by 23 Hz, gray area represent significant windows of interest

#### Position decoding

MUA reliably decoded both representational profiles in both animals, with significant decoding emerging approximately 125 ms after stimulus onset. The decoding of the W-shape peaked at 225 ms in M1 (*R*^2^ = 0.08, *p <* 0.001) and at 125 ms in M2 (*R*^2^ = 0.27, *p <* 0.001). U-shape decoding peaked at a later latency in both monkeys, consistent with the temporal dissociation between early boundary detection and late surface filling-in. Low-frequency LFP activity (5–25 Hz) also reliably decoded both profiles in both animals, though with slightly different preferred frequencies. In M1, W-shape decoding was strongest at 9 Hz (*R*^2^ = 0.04, *p <* 0.001), peaking around 100 ms, whereas U-shape decoding was most robust at 25 Hz (*R*^2^ = 0.11, *p <* 0.001), peaking around 350 ms. In M2, both profiles were best decoded at 10 Hz (W-shape: peak at 100 ms, *R*^2^ = 0.039, *p <* 0.01; U-shape: peak at 175 ms, *R*^2^ = 0.029, *p <* 0.01). Decoding at 25 Hz was weaker in M2 (*R*^2^ = 0.037) and restricted to an early window of 75–100 ms, likely reflecting the initial visual onset response. To maintain consistency with the group-level analyses, 8 Hz was used as the frequency of interest for individual results; this should be noted as a possible source of attenuation for M2, where this frequency was not independently significant for the W-shape.

Temporal generalization of MUA showed partly overlapping patterns across monkeys. In M1, early responses around 100 ms partially generalized to later time points for both shapes; at 8 Hz, generalization was significant but temporally restricted (W-shape: 75–150 ms; U-shape: 175–200 ms). In M2, early MUA responses did not generalize to later time points and stabilized much later; at 8 Hz, no significant generalization was observed for the W-shape, while U-shape generalization was limited to 175–225 ms.

Position–amplitude correlations were significant in M1 (early window: *r* = *−*0.81, *p <* 0.01; later window: *r* = *−*0.63, *p <* 0.01), but not in M2 (early: *r* = *−*0.06, *p* = 0.82; late: *r* = *−*0.37, *p* = 0.28). The absence of significance in M2 likely reflects reduced statistical power due to the smaller number of positions (17 vs. 23).

Figure-ground modulation was strong in both animals, with a higher figure-related maximum in M1 (*FGM_max_*= 0.59) than in M2 (*FGM_max_*= 0.43). Edge-related modulation reached comparable maxima in both animals (M1: *FGM_max_*= 0.53; M2: *FGM_max_* = 0.55).

#### Orientation decoding

Orientation decoding accuracy at IN positions was lower in M2 (*accuracy* = 0.76) than in M1 (*accuracy* = 0.82). In the 100–400 ms window, LFP decoding peaked at 6 Hz at 100 ms in M1 (*accuracy* = 0.72), with mixed weight alignment relative to MUA. In M2, decoding peaked at 9 Hz at 100 ms (*accuracy* = 0.58), with weights oriented opposite to MUA. For cross-temporal generalization, 8 Hz was used to match the main results.

#### Amplitude and phase modulation of MUA

In the IN condition, the relationship between 8 Hz amplitude and MUA differed between monkeys. In M1, MUA was significantly higher in low-amplitude trials (Δ = *−*3.55 *×* 10*^−^*^6^; *p* = 0.002), consistent with the group-level result. In M2, however, the effect was in the opposite direction, with higher MUA in high-amplitude trials (Δ = 2.34 *×* 10*^−^*^7^; *p* = 0.002). This discrepancy may reflect the weaker and less reliable figure-ground modulation in M2 noted above.

Phase-dependent modulation of MUA at 8 Hz was consistent across both monkeys and time windows. In the early window (100–200 ms), significant modulation was observed in M1 (*MoI* = 0.002, *p <* 0.001) and M2 (*MoI* = 0.001, *p <* 0.001), and this effect persisted in the later window (325–425 ms) in both animals (M1: *MoI* = 0.001, *p <* 0.001; M2: *MoI* = 0.003, *p <* 0.001). At 25 Hz, results were less consistent: no significant modulation was observed in M1 in the early window (*MoI* = 0.0002, *p* = 1.0), while M2 showed a significant effect (*MoI* = 0.001, *p* = 0.004); in the later window, neither monkey reached significance after correction (M1: *MoI* = 0.0003, *p* = 1.0; M2: *MoI* = 0.002, *p* = 0.10).

In the OUT condition, 8 Hz phase modulation remained significant in the early window in both monkeys (M1: *MoI* = 0.006, *p <* 0.001; M2: *MoI* = 0.002, *p <* 0.001). In the later window, the effect persisted in M2 (*MoI* = 0.010, *p* = 0.002) but not in M1 (*MoI* = 0.001, *p* = 0.89). No significant phase modulation was observed at 25 Hz in either monkey or time window (early: M1 *MoI* = 0.0004, *p* = 0.719; M2 *MoI* = 0.0007, *p* = 0.348; late: M1 *MoI* = 0.0018, *p* = 0.573; M2 *MoI* = 0.0025, *p* = 0.530).

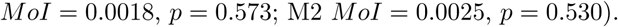

**Figure S7:**
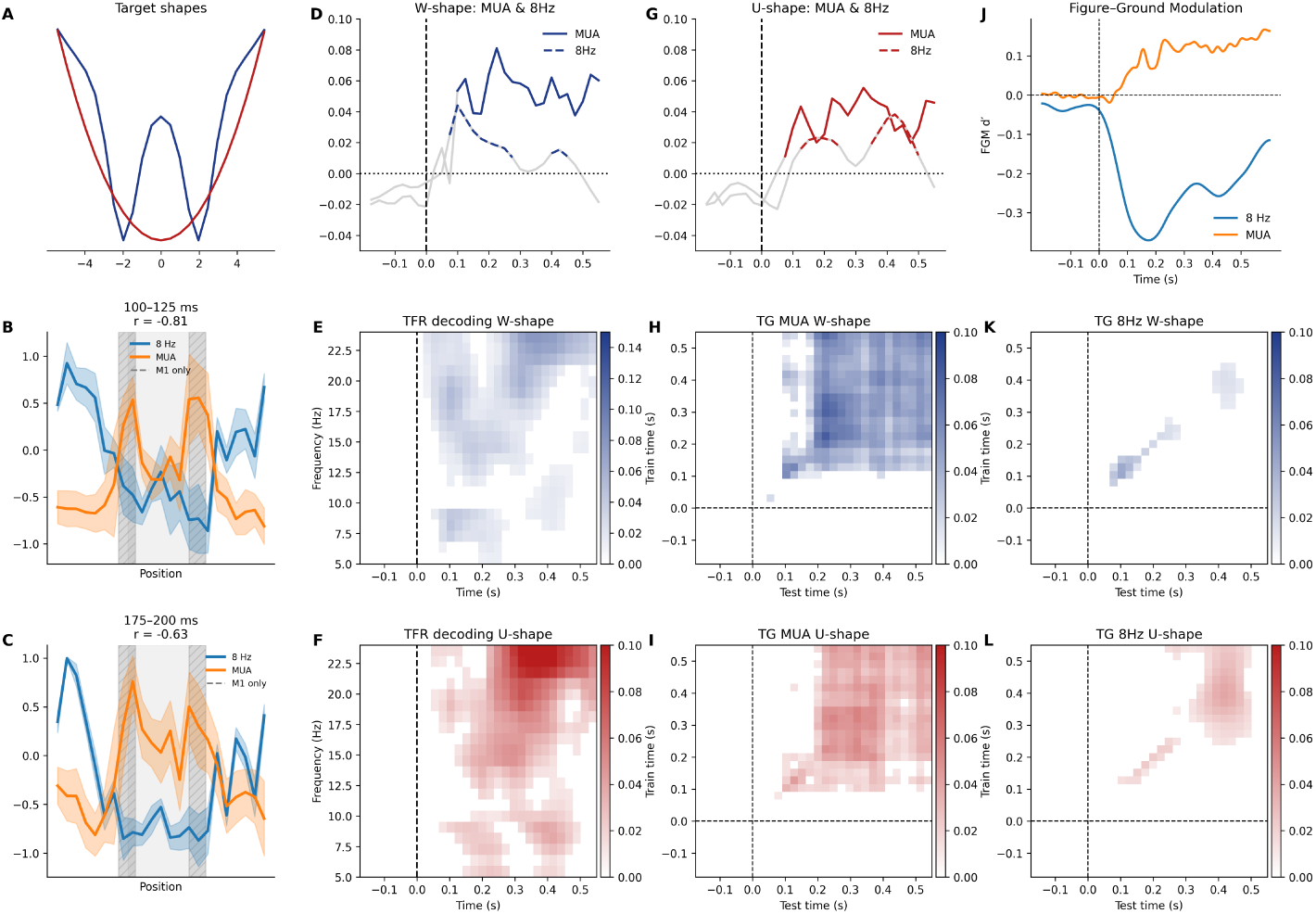
(A) Two testing conditions: blue represent W-shape reflecting edge and center modulation; red represent U-shape reflecting figure filling. (B, C) Averaged MUA and 8 Hz by positions at time windows 100-125 and 175-200 ms (D, G) Decoding results for MUA and LFP 8 Hz for W-shape (blue) and U-shape (red). Solid line MUA, dashed line 8 Hz. (E, F) Decoding results for LFP 5-25 Hz for W-shape (blue) and U-shape (red), colored areas are significant. (H, I) Time generalization of decoding for MUA for W-shape (blue) and U-shape (red), colored areas are significant. (J) Figure-ground modulation orange line MUA, blue line amplitude 8 Hz (K, L) Time generalization of decoding for 8 Hz for W-shape (blue) and U-shape (red), colored areas are significant.

#### V1-V4 interaction

In both monkeys, MI between V1 and V4 MUA was significantly higher in the IN condition than the OUT condition during the 100–200 ms window M1 (peak *MI* = 0.65 at 150 ms) and M2 (peak *MI* = 0.18 at 175 ms), consistent with the weaker figure-ground modulation in M2.

Amplitude-dependent modulation of inter-areal MI showed a consistent pattern in V1: MI was higher in low-alpha trials in both monkeys and both conditions. In the IN condition, V1 effects were significant in M1 (Δ*MI* = *−*0.195; *p*0.002) and M2 (Δ*MI* = *−*0.060; *p* = 0.002). In the OUT condition, the same direction was observed in M1 (Δ*MI* = *−*0.171; *p* = 0.002) and M2 (Δ*MI* = *−*0.058; *p* = 0.002).

In V4, the pattern was asymmetric and less consistent across monkeys. In the IN condition, MI was higher in *high*-alpha trials in M1 (Δ*MI* = +0.066; *p* = 0.019), while the effect in M2 was not significant (Δ*MI* = +0.012; *p* = 0.278). In the OUT condition, both monkeys showed higher MI in low-alpha trials, though only M1 reached significance (M1: Δ*MI* = *−*0.134, *p* = 0.012; M2: Δ*MI* = *−*0.030, *p* = 0.086). The reversal of the V4 effect between IN and OUT conditions in M1 is consistent with the group-level result. Phase-dependent modulation of inter-areal MI is reported in the main text.

**Figure S8:**
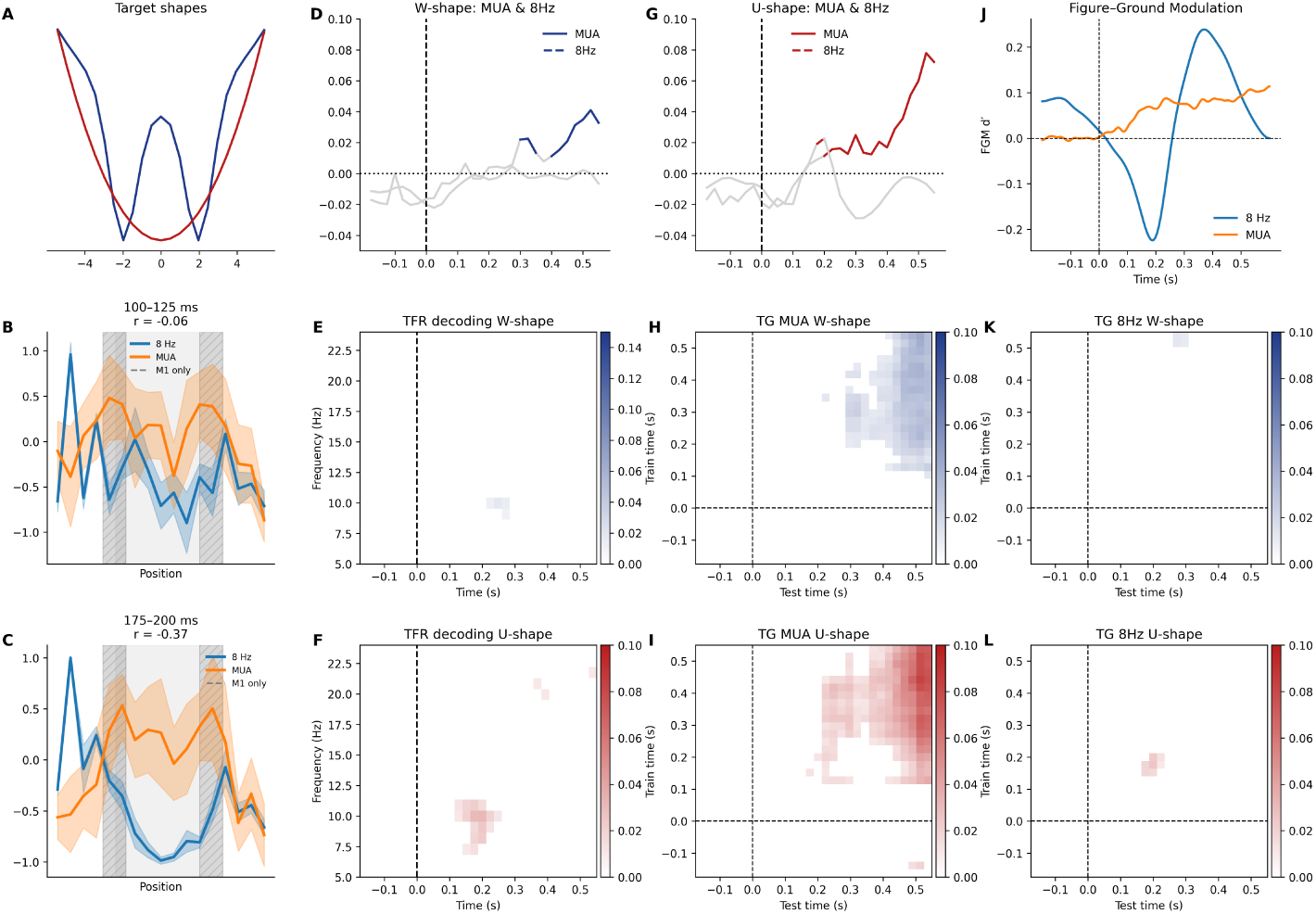
(A) Two testing conditions: blue represent W-shape reflecting edge and center modulation; red represent U-shape reflecting figure filling. (B, C) Averaged MUA and 8 Hz by positions at time windows 100-125 and 175-200 ms (D, G) Decoding results for MUA and LFP 8 Hz for W-shape (blue) and U-shape (red). Solid line MUA, dashed line 8 Hz. (E, F) Decoding results for LFP 5-25 Hz for W-shape (blue) and U-shape (red), colored areas are significant. (H, I) Time generalization of decoding for MUA for W-shape (blue) and U-shape (red), colored areas are significant. (J) Figure-ground modulation orange line MUA, blue line amplitude 8 Hz (K, L) Time generalization of decoding for 8 Hz for W-shape (blue) and U-shape (red), colored areas are significant.

#### Orientation-selective channels

The difference in spectral power between preferred and non-preferred orientations revealed distinct frequency-specific dynamics in the two monkeys. In M1, a positive peak was observed at approximately 7 Hz around 90 ms after stimulus onset, followed by a negative peak near 14 Hz at approximately 135 ms. In contrast, M2 exhibited an early positive peak at around 20 Hz near 60 ms, followed by a negative peak centered at approximately 6 Hz around 100 ms.

**Figure S9:**
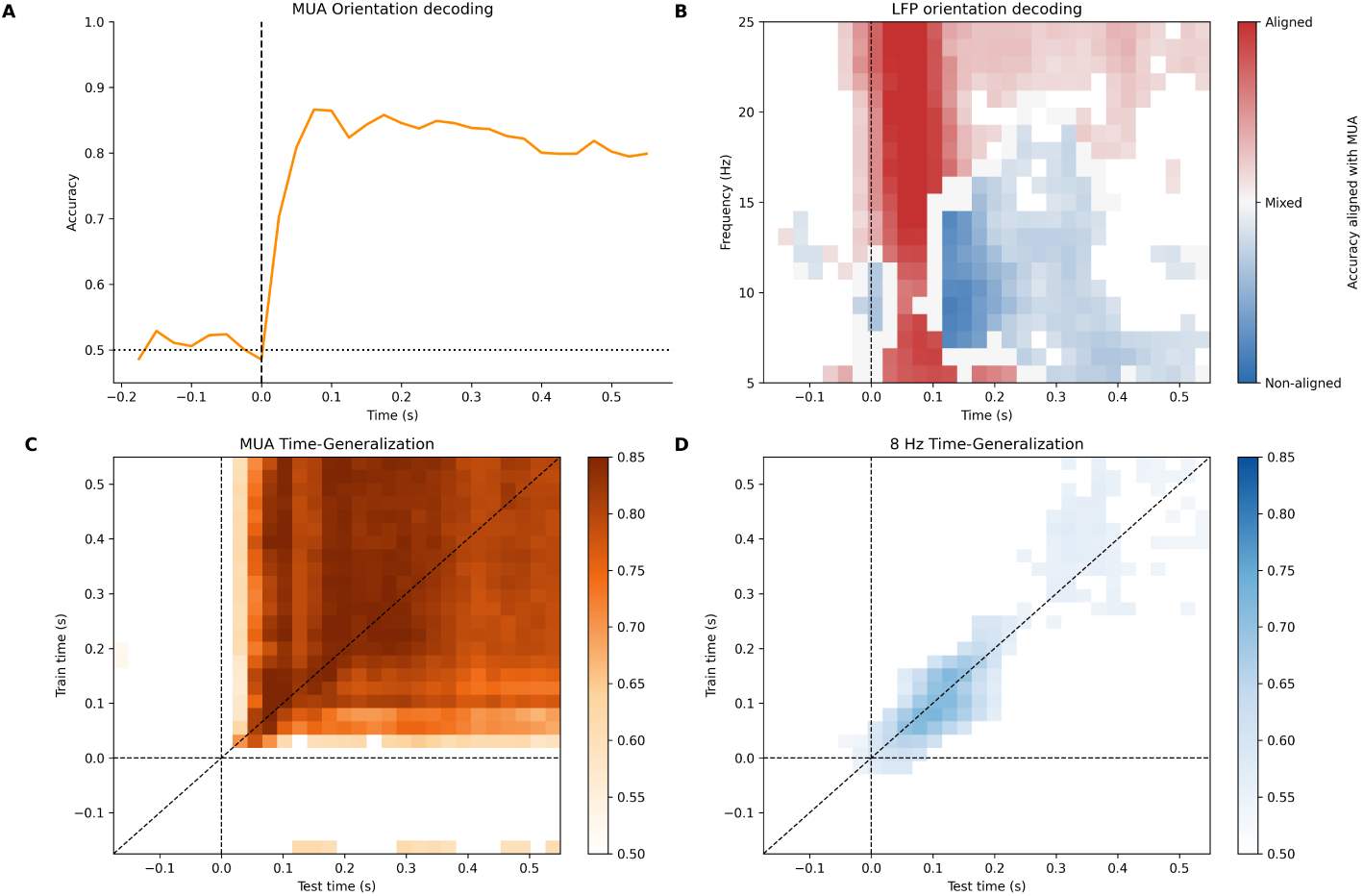
Orientation decoding at IN positions for M1. (A) MUA decoding accuracy over time. (B) LFP decoding: grey regions indicate significant decoding with no consistent weight alignment; blue regions indicate weights opposing MUA; red regions indicate weights aligned with MUA. Colour intensity reflects normalised accuracy (0.50–0.75). (C) MUA cross-temporal generalisation. (D) 8 Hz LFP cross-temporal generalisation.

Orientation-selective channels showed higher 13 Hz amplitude than non-selective channels in M1 (Δ = 93, *p <* 0.001), but not in M2 (Δ = *−*1, *p* = 0.74). Amplitude-dependent MUA modulation at 13 Hz was significant in both monkeys (M1: Δ = 5.37 *×* 10*^−^*^7^, *p* = 0.007; M2: Δ = 4.09 *×* 10*^−^*^7^, *p* = 0.02). Phase–MUA coupling at 13 Hz was not significant in either monkey after FDR correction (M1: *MoI* = 0.0002, *p* = 0.842; M2: *MoI* = 0.0003, *p* = 0.376). Inter-areal MI did not differ significantly between high- and low-13 Hz-amplitude groups (M1: Δ*MI* = *−*0.01, *p* = 0.03; M2: Δ*MI* = *−*0.0007, *p* = 0.089).

**Figure S10:**
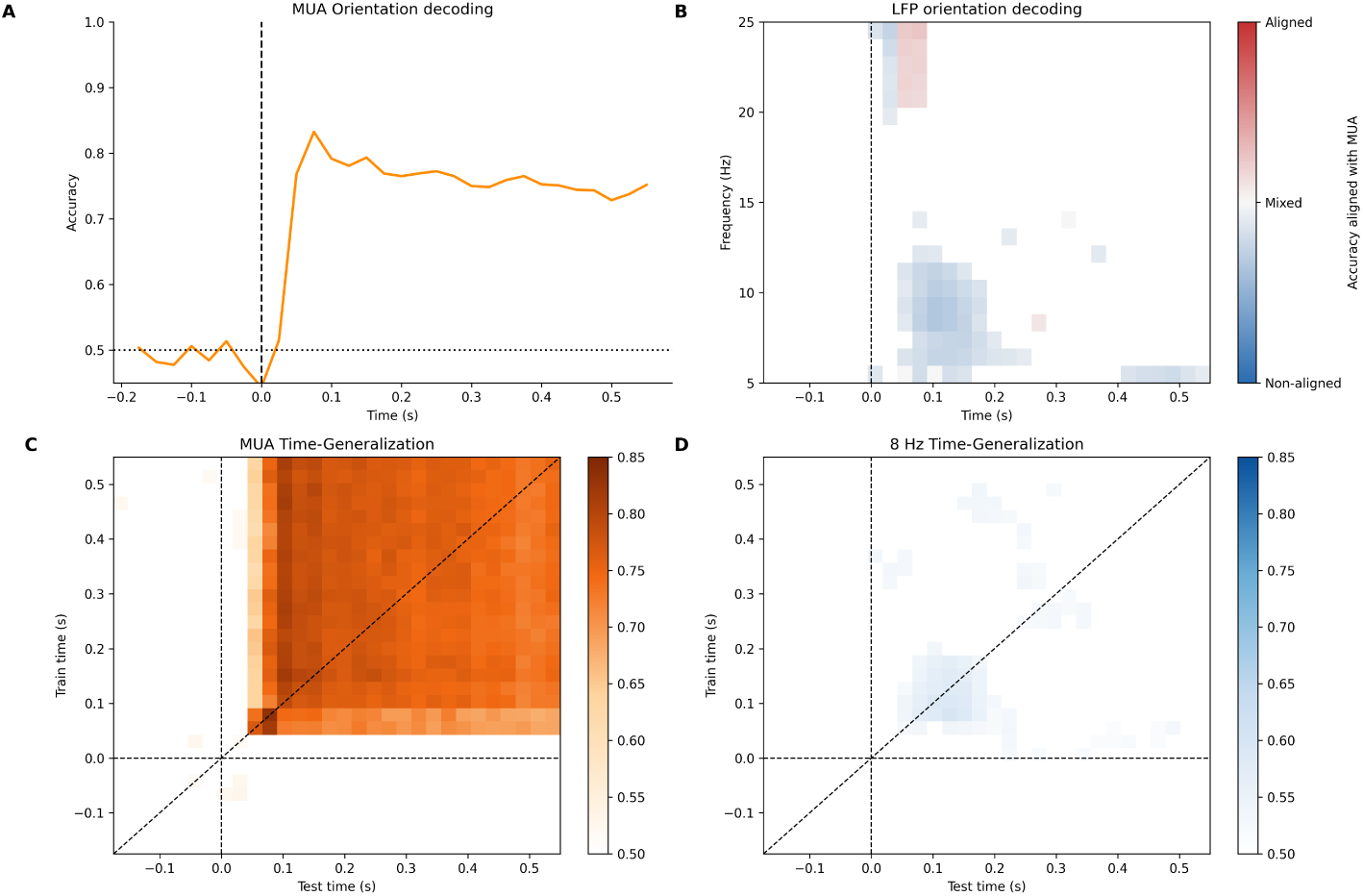
Orientation decoding at IN positions for M2. (A) MUA decoding accuracy over time. (B) LFP decoding: grey regions indicate significant decoding with no consistent weight alignment; blue regions indicate weights opposing MUA; red regions indicate weights aligned with MUA. Colour intensity reflects normalised accuracy (0.50–0.75). (C) MUA cross-temporal generalisation. (D) 8 Hz LFP cross-temporal generalisation.

**Figure S11:**
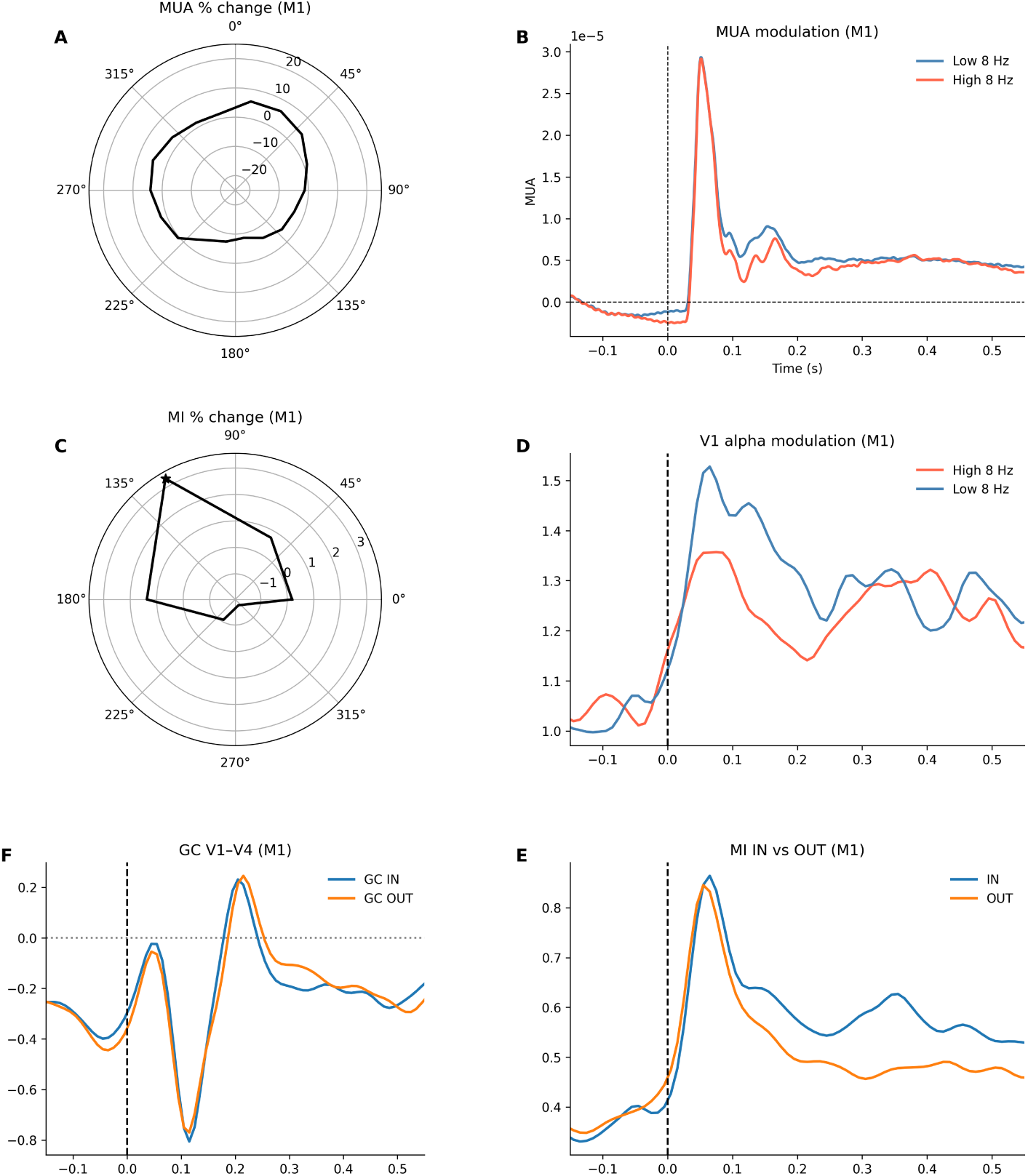
V1-V4 interaction results for M1. (A) MUA modulated by 8 Hz phase in the 100–200 ms window. (B) MUA in low-(blue) and high-(orange) amplitude 8 Hz trials; grey region indicates the significant window. (C) Inter-areal MI of V1–V4 phase difference at 8 Hz, expressed as percentage change from the bin mean; solid line M1, dashed line M2. (D) MI in the IN condition for low-(orange) and high-(black) alpha trials. (E) MI in IN (blue) and OUT (orange) conditions; grey region indicates the significant window. (F) Granger causality over time at 8 Hz: positive values indicate V1*→*V4 dominance, negative values indicate V4*→*V1 dominance.

**Figure S12:**
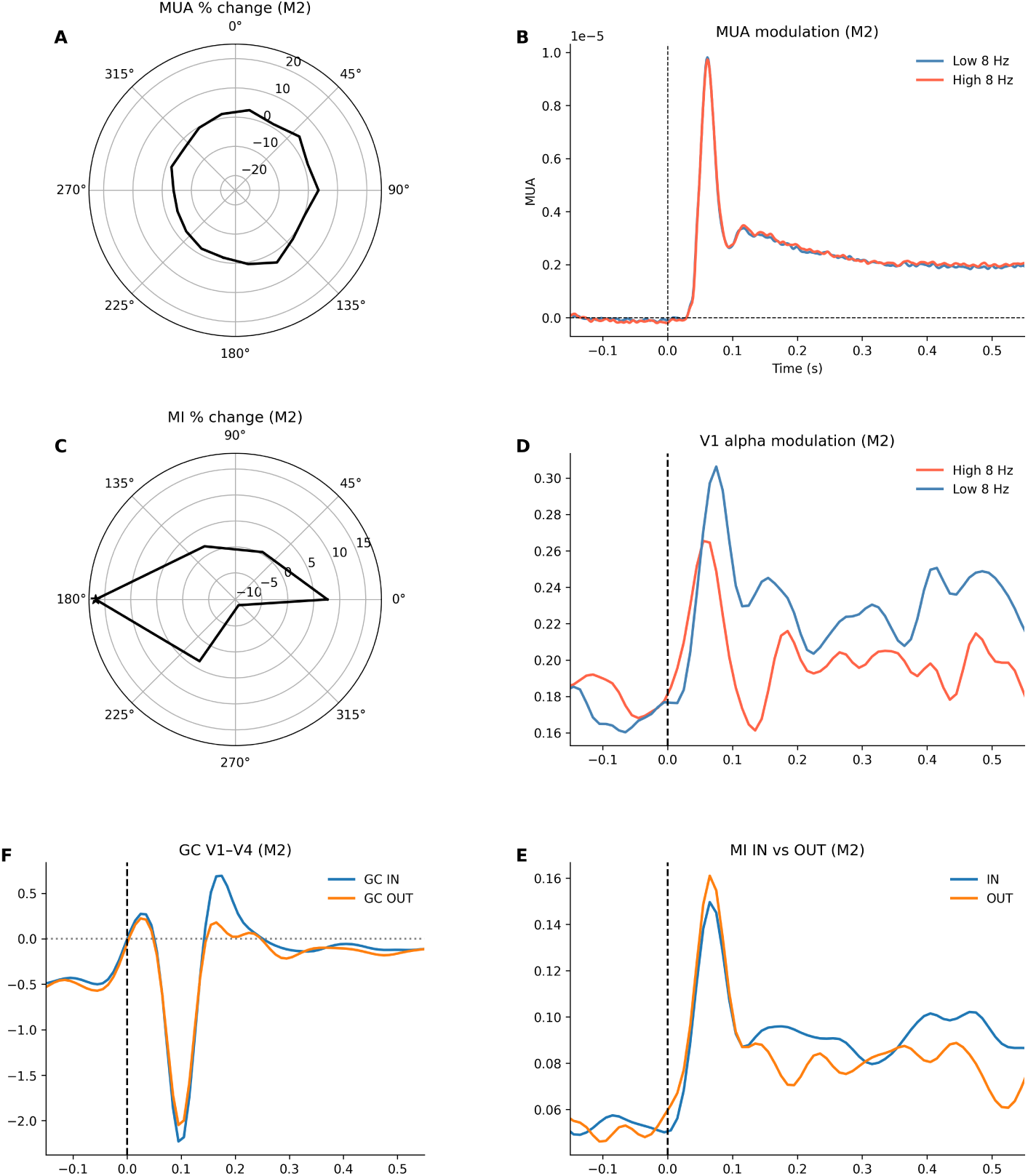
V1-V4 interaction results for M2. (A) MUA modulated by 8 Hz phase in the 100–200 ms window. (B) MUA in low-(blue) and high-(orange) amplitude 8 Hz trials; grey region indicates the significant window. (C) Inter-areal MI of V1–V4 phase difference at 8 Hz, expressed as percentage change from the bin mean; solid line M1, dashed line M2. (D) MI in the IN condition for low-(orange) and high-(black) alpha trials. (E) MI in IN (blue) and OUT (orange) conditions; grey region indicates the significant window. (F) Granger causality over time at 8 Hz: positive values indicate V1*→*V4 dominance, negative values indicate V4*→*V1 dominance.

**Figure S13:**
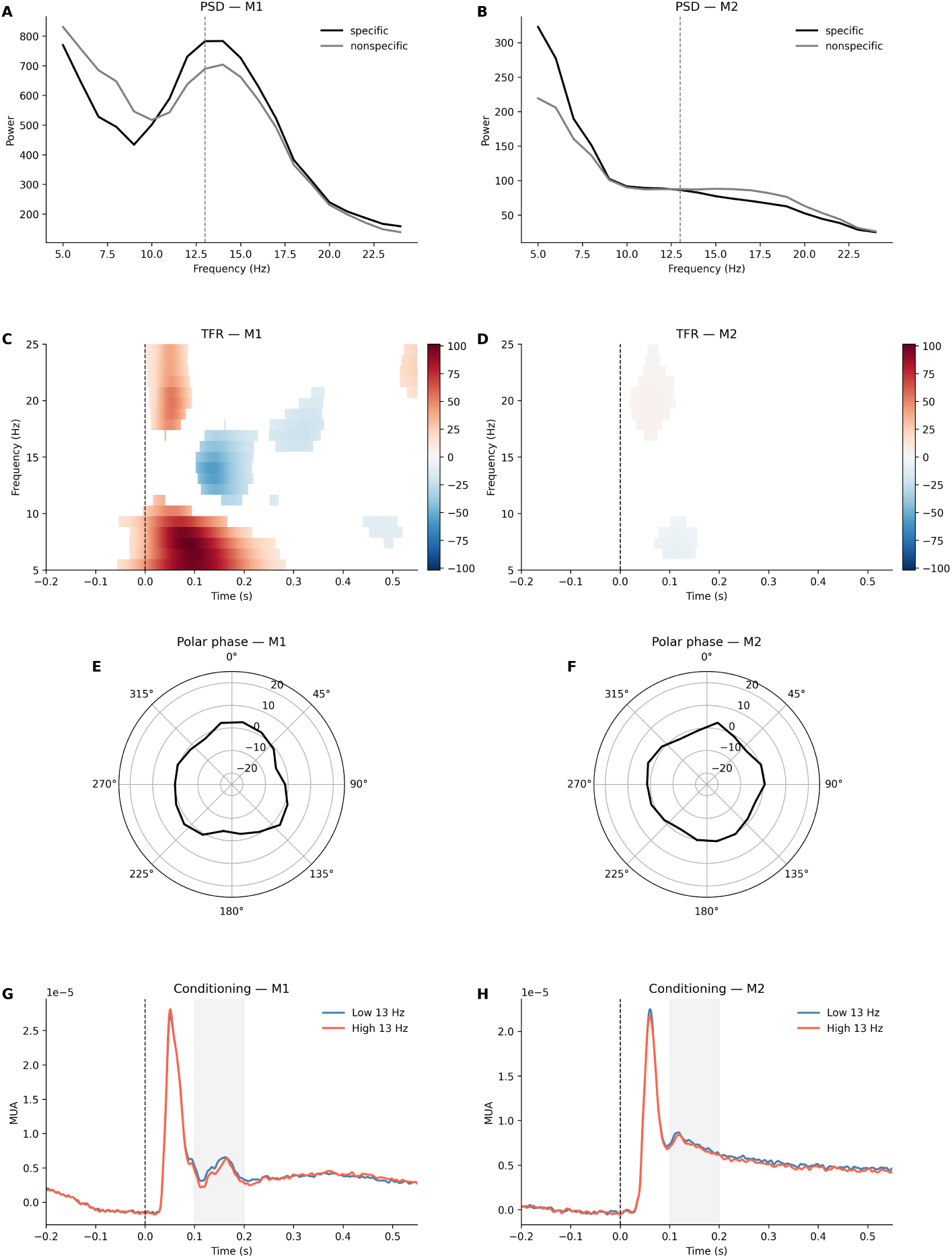
(A/B) Power spectral density for channels with a preferred orientation (blue solidline) and channels without a preferred orientation (blue dashed line) (C/D) The difference between the preferred and non-preferred orientations for frequency amplitudes from 5 to 25. Significant windows are marked in color, insignificant ones in white. (E/F) MUA phasse modulation by 13 Hz (G/H) MUA amplitude modulation by 13 Hz, gray area represent significant window of interest

## References

Bastos, A. M., Vezoli, J., Bosman, C. A., Schoffelen, J.-M., Oostenveld, R., Dowdall, J. R., De Weerd, P., Kennedy, H., & Fries, P. (2015). Visual areas exert feedforward and feedback influences through distinct frequency channels. Neuron, 85 (2), 390–401. 10.1016/j.neuron.2014.12.018

Bollimunta, A., Chen, Y., Schroeder, C. E., & Ding, M. (2008). Neuronal mechanisms of cortical alpha oscillations in awake-behaving macaques. Journal of Neuroscience, 28 (40), 9976–9988. 10.1523/JNEUROSCI.2300-08.2008

Bollimunta, A., Mo, J., Schroeder, C. E., & Ding, M. (2011). Neuronal mechanisms and attentional modulation of corticothalamic alpha oscillations. Journal of Neuroscience, 31 (13), 4935–4943. 10.1523/JNEUROSCI.5580-10.2011

Bonnefond, M., & Jensen, O. (2015). Gamma activity coupled to alpha phase as a mechanism for top-down controlled gating. PLOS ONE, 10 (6), e0128667. 10.1371/journal.pone.0128667

Bonnefond, M., Kastner, S., & Jensen, O. (2017). Communication between brain areas based on nested oscillations. eNeuro, 4 (2), ENEURO.0153–16.2017. 10.1523/ENEURO.0153-16.2017

Buzsáki, G., & Draguhn, A. (2004). Neuronal oscillations in cortical networks. Science, 304 (5679), 1926–1929. 10.1126/science.1099745

Clausner, T., Marques, J. P., Scheeringa, R., & Bonnefond, M. (2025). Frequency and laminar profile of feature specific visual activity revealed by inter-leaved EEG-fMRI. eLife, 14. 10.7554/eLife.108408.1

Clayton, M. S., Yeung, N., & Cohen Kadosh, R. (2018). The many characters of visual alpha oscillations. European Journal of Neuroscience, 48 (7), 2498–2508. 10.1111/ejn.13747

Di Dona, G., & Ronconi, L. (2023). Beta oscillations in vision: A (preconscious) neural mechanism for the dorsal visual stream? Frontiers in Psychology, 14, 1296483. 10.3389/fpsyg.2023.1296483

Ding, Z., Tran, D., Ponder, K., Ding, Z., Froebe, R., Ntanavara, L., Fahey, P. G., Cobos, E., Baroni, L., Diamantaki, M., Wang, E. Y., Chang, A., Papadopoulos, S., Fu, J., Muhammad, T., Papadopoulos, C., Cadena, S. A., Evangelou, A., Willeke, K., … Tolias, A. S. (2026). Functional bipartite invariance in mouse primary visual cortex receptive fields. Nature Neuroscience, 29, 851–863. 10.1038/s41593-026-02213-3

Dougherty, K., Cox, M. A., Ninomiya, T., Leopold, D. A., & Maier, A. (2017). Ongoing alpha activity in visual cortex inhibits spiking responses to oriented stimuli. Journal of Neuroscience, 37 (14), 3700–3713. 10.1523/JNEUROSCI.2330-16.2017

Engel, A. K., & Fries, P. (2010). Beta-band oscillations — signalling the status quo? Current Opinion in Neurobiology, 20 (2), 156–165. 10.1016/j.conb.2010.02.015

Fiebelkorn, I. C., Pinsk, M. A., & Kastner, S. (2018). A dynamic interplay within the frontoparietal network underlies rhythmic spatial attention. Neuron, 99 (4), 842–853. 10.1016/j.neuron.2018.07.038

Fries, P. (2005). A mechanism for cognitive dynamics: Neuronal communication through neuronal coherence. Trends in Cognitive Sciences, 9 (10), 474–480. 10.1016/j.tics.2005.08.011

Fries, P. (2015). Rhythms for cognition: Communication through coherence. Neuron, 88 (1), 220–235. 10.1016/j.neuron.2015.09.034

Gabhart, K. M., Xiong, Y. S., & Bastos, A. M. (2025). Predictive coding: A more cognitive process than we thought? Trends in Cognitive Sciences. 10.1016/j.tics.2025.01.012

Harvey, B. M., & Dumoulin, S. O. (2013). The relationship between cortical magnification factor and population receptive field size in human visual cortex: Constancies in cortical architecture. Journal of Neuroscience, 33 (6), 2317–2325. 10.1523/JNEUROSCI.2572-12.2013

Haufe, S., Meinecke, F., Görgen, K., Dähne, S., Haynes, J.-D., Blankertz, B., & Bießmann, F. (2014). On the interpretation of weight vectors of linear models in multivariate neuroimaging. NeuroImage, 87, 96–110. 10.1016/j.neuroimage.2013.10.067

Hoppensteadt, F. C., & Izhikevich, E. M. (1998). Thalamocortical interactions modeled by weakly connected oscillators: Could the brain use FM radio principles? Biosystems, 48 (1–3), 85–94. 10.1016/S0303-2647(98)00053-7

Huang, X., & Paradiso, M. A. (2008). V1 response timing and surface filling-in. Journal of Neurophysiology, 100 (1), 539–547. 10.1152/jn.90397.2008

Ince, R. A. A., Giordano, B. L., Kayser, C., Rousselet, G. A., Gross, J., & Schyns, P. G. (2017). A statistical framework for neuroimaging data analysis based on mutual information estimated via a gaussian copula. NeuroImage, 150, 54–63. 10.1016/j.neuroimage.2017.02.041

Jensen, O., & Bonnefond, M. (2026). The alpha rhythm: From physiology to behaviour. Physiological Reviews. 10.1152/physrev.00001.2025

Jensen, O., & Mazaheri, A. (2010). Shaping functional architecture by oscillatory alpha activity: Gating by inhibition. Frontiers in Human Neuro-science, 4, 186. 10.3389/fnhum.2010.00186

Jeurissen, D., van Ham, A. F., Gilhuis, A., et al. (2024). Border-ownership tuning determines the connectivity between v4 and v1 in the macaque visual system. Nature Communications, 15, 9115. 10.1038/s41467-024-53256-8

Kelly, S. P., Lalor, E. C., Reilly, R. B., & Foxe, J. J. (2006). Increases in alpha oscillatory power reflect an active retinotopic mechanism for distracter suppression during sustained visuospatial attention. Journal of Neuro-physiology, 95 (6), 3844–3851. 10.1152/jn.01234.2005

King, J.-R., & Dehaene, S. (2014). Characterizing the dynamics of mental representations: The temporal generalization method. Trends in Cognitive Sciences, 18 (4), 203–210. 10.1016/j.tics.2014.01.002

Kirchberger, L., Mukherjee, S., Schnabel, U. H., van Beest, E. H., Barsegyan, A., Levelt, C. N., & Roelfsema, P. R. (2021). The essential role of recur-rent processing for figure-ground perception in mice. Science Advances, 7 (27), eabe1833. 10.1126/sciadv.abe1833

Klimesch, W. (2012). Alpha-band oscillations, attention, and controlled access to stored information. Trends in Cognitive Sciences, 16 (12), 606–617. 10.1016/j.tics.2012.10.007

Klimesch, W., Sauseng, P., & Hanslmayr, S. (2007). Eeg alpha oscillations: The inhibition–timing hypothesis. Brain Research Reviews, 53 (1), 63–88. 10.1016/j.brainresrev.2006.06.003

Klink, P. C., Dagnino, B., Gariel-Mathis, M.-A., & Roelfsema, P. R. (2017). Distinct feedforward and feedback effects of microstimulation in visual cortex reveal neural mechanisms of texture segregation. Neuron, 95 (1), 209–220. 10.1016/j.neuron.2017.05.033

Lamme, V. A. F., Zipser, K., & Spekreijse, H. (1998). Figure–ground activity in primary visual cortex is suppressed by anesthesia. Proceedings of the National Academy of Sciences, 95 (6), 3263–3268. 10.1073/pnas.95.6.3263

Markov, N. T., Misery, P., Falchier, A., Lamy, C., Vezoli, J., Quilodran, R., Gariel, M. A., Giroud, P., Ercsey-Ravasz, M., Pilaz, L. J., Huissoud, C., Barone, P., Dehay, C., Toroczkai, Z., Van Essen, D. C., Kennedy, H., & Knoblauch, K. (2011). Weight consistency specifies regularities of macaque cortical networks. Cerebral Cortex, 21 (6), 1254–1272. 10.1093/cercor/bhq201

Poort, J., Raudies, F., Wannig, A., Lamme, V. A. F., Neumann, H., & Roelf-sema, P. R. (2012). The role of attention in figure–ground segregation in areas V1 and V4 of the visual cortex. Neuron, 75 (1), 143–156. 10.1016/j.neuron.2012.04.032

Poort, J., Self, M. W., van Vugt, B., Malkki, H., & Roelfsema, P. R. (2016). Texture segregation causes early figure enhancement and later ground suppression in areas V1 and V4 of visual cortex. Cerebral Cortex, 26 (10), 3964–3976. 10.1093/cercor/bhv186

Samaha, J., Gosseries, O., & Postle, B. R. (2016). Distinct oscillatory frequencies underlie excitability of human occipital and parietal cortex. Journal of Neuroscience, 36 (6), 2093–2102. 10.1523/JNEUROSCI.3rible-15.2016

Self, M. W., Jeurissen, D., van Ham, A. F., van Vugt, B., Poort, J., & Roelfsema, P. R. (2019). The segmentation of proto-objects in the monkey primary visual cortex. Current Biology, 29 (6), 1019–1029. 10.1016/j.cub.2019.02.016

Self, M. W., van Kerkoerle, T., Supèr, H., & Roelfsema, P. R. (2013). Distinct roles of the cortical layers of area V1 in figure–ground segregation. Cur-rent Biology, 23 (21), 2121–2129. 10.1016/j.cub.2013.09.013

Seymour, R. A., Rippon, G., Gooding-Williams, G., Schoffelen, J. M., & Kessler, K. (2019). Dysregulated oscillatory connectivity in the visual system in autism spectrum disorder. Brain, 142 (10), 3294–3305. 10.1093/brain/awz214

Singer, W. (2018). Neuronal oscillations: Unavoidable and useful? European Journal of Neuroscience, 48 (7), 2389–2398. 10.1111/ejn.14088

Singer, W., & Effenberger, F. (2025). Oscillations in natural neuronal networks: An epiphenomenon or a fundamental computational mechanism? Hu-man Arenas. 10.1007/s42087-025-00478-x

Spaak, E., Bonnefond, M., Maier, A., Leopold, D. A., & Jensen, O. (2012). Layer-specific entrainment of gamma-band neural activity by the alpha rhythm in monkey visual cortex. Current Biology, 22 (24), 2313–2318. 10.1016/j.cub.2012.10.020

Supèr, H., Spekreijse, H., & Lamme, V. A. F. (2001). A neural correlate of working memory in the monkey primary visual cortex. Science, 293 (5527), 120–124. 10.1126/science.1060496

Tort, A. B. L., Komorowski, R., Eichenbaum, H., & Kopell, N. (2010). Measuring phase-amplitude coupling between neuronal oscillations of different frequencies. Journal of Neurophysiology, 104 (2), 1195–1210. 10.1152/jn.00106.2010

van Kerkoerle, T., Self, M. W., Dagnino, B., Gariel-Mathis, M.-A., Poort, J., van der Togt, C., & Roelfsema, P. R. (2014). Alpha and gamma oscillations characterize feedback and feedforward processing in monkey visual cortex. Proceedings of the National Academy of Sciences, 111 (40), 14332–14341. 10.1073/pnas.1402773111

Wen, X., Chang, Y., Li, S., Wang, J., Li, X., Li, D., & Liang, Z. (2025). A practical measure of integrated information reveals alpha-band activity and the posterior cortex as neural correlates of arousal. NeuroImage, 121384. 10.1016/j.neuroimage.2025.121384

Westerberg, J., & Roelfsema, P. (2025). Hierarchical interactions between sensory cortices defy predictive coding. Trends in Cognitive Sciences, 30, 110–123. 10.1016/j.tics.2025.09.018

Yuasa, K., Groen, I. I., Piantoni, G., Montenegro, S., Flinker, A., Devore, S., & Winawer, J. (2025). Precise spatial tuning of visually driven alpha oscillations in human visual cortex. eLife, 12, RP90387. 10.7554/eLife.90387

